# OME-Zarr: a cloud-optimized bioimaging file format with international community support

**DOI:** 10.1101/2023.02.17.528834

**Authors:** Josh Moore, Daniela Basurto-Lozada, Sébastien Besson, John Bogovic, Jordão Bragantini, Eva M. Brown, Jean-Marie Burel, Xavier Casas Moreno, Gustavo de Medeiros, Erin E. Diel, David Gault, Satrajit S. Ghosh, Ilan Gold, Yaroslav O. Halchenko, Matthew Hartley, Dave Horsfall, Mark S. Keller, Mark Kittisopikul, Gabor Kovacs, Aybüke Küpcü Yoldaş, Koji Kyoda, Albane le Tournoulx de la Villegeorges, Tong Li, Prisca Liberali, Dominik Lindner, Melissa Linkert, Joel Lüthi, Jeremy Maitin-Shepard, Trevor Manz, Luca Marconato, Matthew McCormick, Merlin Lange, Khaled Mohamed, William Moore, Nils Norlin, Wei Ouyang, Bugra Özdemir, Giovanni Palla, Constantin Pape, Lucas Pelkmans, Tobias Pietzsch, Stephan Preibisch, Martin Prete, Norman Rzepka, Sameeul Samee, Nicholas Schaub, Hythem Sidky, Ahmet Can Solak, David R. Stirling, Jonathan Striebel, Christian Tischer, Daniel Toloudis, Isaac Virshup, Petr Walczysko, Alan M. Watson, Erin Weisbart, Frances Wong, Kevin A. Yamauchi, Omer Bayraktar, Beth A. Cimini, Nils Gehlenborg, Muzlifah Haniffa, Nathan Hotaling, Shuichi Onami, Loic A. Royer, Stephan Saalfeld, Oliver Stegle, Fabian J. Theis, Jason R. Swedlow

## Abstract

A growing community is constructing a next-generation file format (NGFF) for bioimaging to overcome problems of scalability and heterogeneity. Organized by the Open Microscopy Environment (OME), individuals and institutes across diverse modalities facing these problems have designed a format specification process (OME-NGFF) to address these needs. This paper brings together a wide range of those community members to describe the cloud-optimized format itself – OME-Zarr – along with tools and data resources available today to increase FAIR access and remove barriers in the scientific process. The current momentum offers an opportunity to unify a key component of the bioimaging domain — the file format that underlies so many personal, institutional, and global data management and analysis tasks.

## Introduction

The exchange of scientific data is one of the key hallmarks of scientific practice in the 21^st^ century. In 2016 Wilkinson and colleagues provided guidelines for making scientific data findable, accessible, interoperable, and reusable (FAIR) that provide a foundation for future scientific discoveries through data integration, reanalysis and the development of new analytic tools (Wilkinson et al. 2016). In the case of biological and biomedical imaging (collectively, “bioimaging”), the size, complexity and heterogeneity of datasets present several impediments towards that goal, the most immediate of which are the specification and construction of data formats that can meet the requirements of FAIR data (Könnecke et al. 2015).

Any format must support both the pixel measurements that are the core of bioimaging data as well as relevant imaging metadata. Specifications that enable storage of experimental, acquisition, and analytic metadata are necessary. The implementation of metadata specifications must be both flexible and scalable to handle the large and heterogeneous volumes of analytic metadata generated, for example the definition of the segmentations and annotations on individual cells and tissues that are quite common in biological imaging workflows. Critically, the set of formats available to end users must support local data storage (laptops, desktop computers, etc.) as well as cloud-based storage that is becoming more heavily used as dataset volumes grow.

Previously, the Open Microscopy Environment (OME) developed OME-TIFF as an open-source file format in bioimaging. Accompanied by reference software implementations, OME-TIFF is primarily for use in fluorescence imaging workflows and has recently been updated to enable whole slide imaging technologies (Besson et al. 2019). This format combines the fundamentally 2D TIFF format with metadata cast in XML in the TIFF header. Its structure makes it appropriate for many applications, where the plane-based access pattern is appropriate.

For bioimaging applications that require large non-planar access to volume data, e.g., arbitrary slicing from user-defined angles, a more sophisticated “chunking” of the data is required that defines how data is stored in accessible and reasonable subsections. This means that large, multi-Gigabyte up to Petabyte bioimaging datasets are not accessed all at once but can be accessed in reasonably sized planes or sub-volumes. In the case of TIFF, the chunk is a tile of the 2D plane allowing data access across time-lapse series or 3D volume. N-dimensional formats like HDF5 (“Hierarchical Data Format”) provide much more flexibility and allow chunking across different dimensions chosen by the user. While TIFF and HDF5 are well established, the chunking strategies depend on fast random access to the entire file that is common in laptops, desktop machines and large cluster file systems, but is not provided by large scalable cloud-based object storage.

Over the last few years, a new data format, Zarr^1^, has been developed for the storage of large N-dimensional typed arrays in the cloud. The Zarr format is now heavily adopted across many scientific communities from genomics to astrophysics (Miles et al. 2023). Zarr stores associated metadata in JSON and binary data in individually referenceable “chunk”-files, providing a flexible, scalable method for storing multidimensional data. In 2021, OME published the first specification and example uses of a “next-generation file format” (NGFF) in bioimaging using the Zarr format (Moore et al. 2021). The first versions of this format, OME-Zarr, focused on developing functionality that tests and demonstrates the utility of the format in bioimaging domains that routinely generate large, metadata-rich datasets – high content screening, digital pathology, electron microscopy, and light sheet imaging.

The discussions necessary to arrive at these specifications have also presented an opportunity to build a coherent development community under the OME-NGFF umbrella, combining a growing range of use cases and requirements with an open, transparent, but technically valid development process. The result has been a thriving community based on open development and open-source principles (Rueden et al. 2019). This open, collaborative approach has been essential to tackle the addition of complex additional metadata to OME-Zarr. This was important as neither TIFF nor HDF5 has specifications for many of the derived data types that are generated in an analysis workflow, e.g., regions of interests (ROIs), labels and other derived data which are crucial in modern analysis workflows. In most cases accessory files are generated to handle these limitations but as data volumes grow, these create additional problems for management and linkage of data. Using the established development process, this functionality was first formally adopted into the OME-NGFF specification, then added to the OME-Zarr implementation, but can equally be applied to other formats like HDF5 in the future.

In this paper, we review the current status of the OME-Zarr format and focus on resources that are now available to users for creating, accessing, and visualizing data stored in OME-Zarr (Fig. 1). This report is timely as we have seen a rapid expansion in tools that support OME-Zarr since the first publication. We also report on the growth of adoption of OME-Zarr in public data repositories. This survey by active members of the OME-NGFF community is meant to provide an update on the status of the ecosystem that has grown around the format and the development community that is developing and releasing tools that can be used by the broader bioimaging community.

**Fig. 1.**
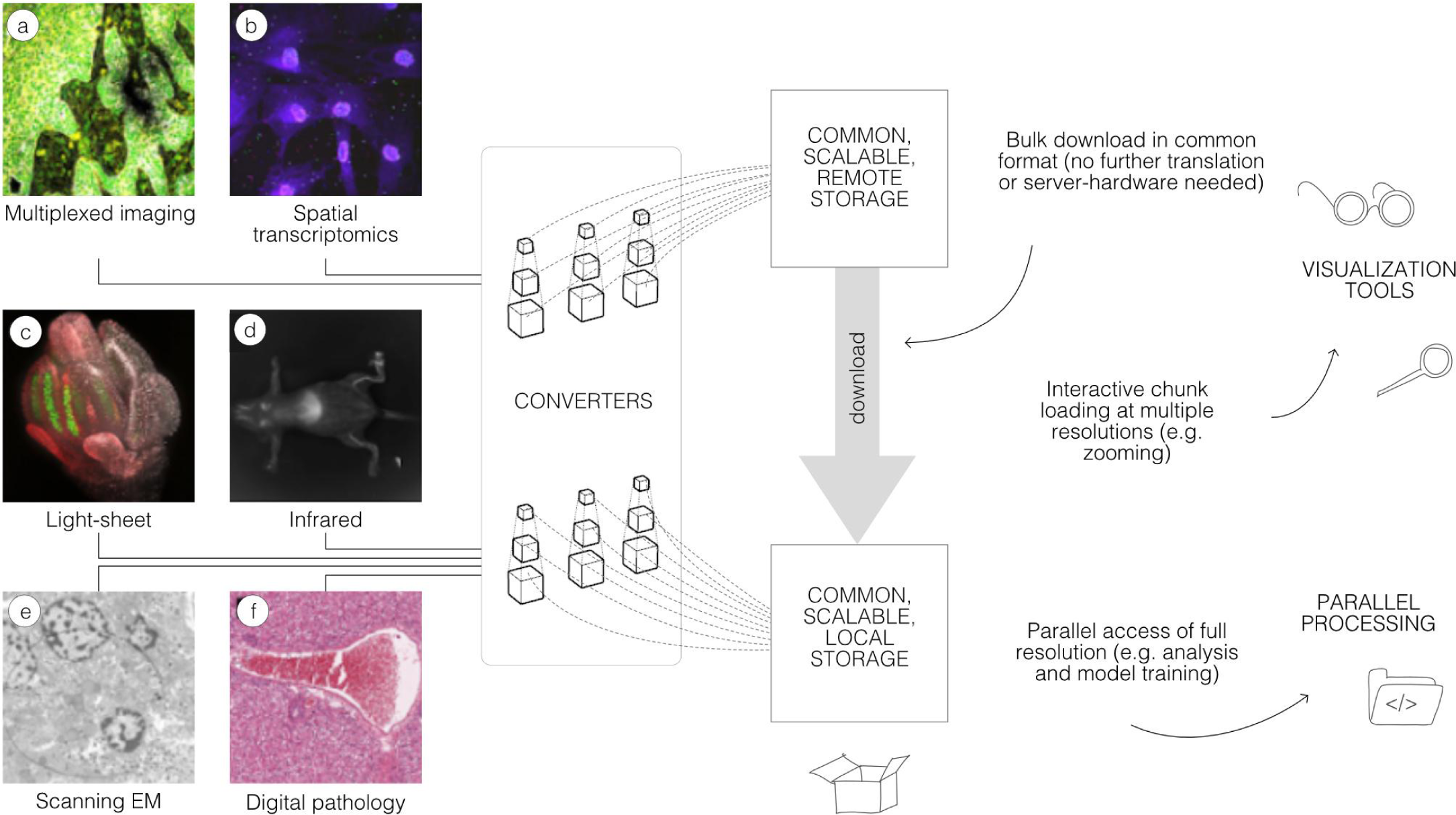
A common format enables a diverse set of use cases via a consistent API. A wide range of modalities can be converted into a representation that can be equally accessed by a variety of tools. This format can be used to download entire datasets for local processing, to stream pyramidal sub-resolutions for interactive viewing or to process entire r solutions in parallel. OME-Zarr data shown includes (A) idr0076 (Ali et al. 2020), (B) idr0101 (Payne et al. 2021), (C) idr0077 (Valuchova et al. 2020), (D) S-BIAD548 (Lim et al. 2023), (E) S-BIAD217 (de Boer et al. 2020), and (F) S-BIAD501 (Igarashi et al. 2015).

## Growth of a compatible solution

The development of a common format is not a light undertaking. Historical approaches to address challenges of scale most often offer a problem-specific and highly-optimized solution, and do not generalize to the wider bioimaging community, reducing interoperability and re-use of datasets and software. Bespoke formats are often incompatible and require significant time and compute resources spent in data wrangling, and generally reduce the amount of FAIR data that is available to scientists. Without a formal body to declare such specifications or dedicated funding to produce a single solution for users, work is left to the community to discuss and implement with the available resources. The larger the community consensus, the more tools can be adapted with the agreed upon solution. In turn, the lives of the users in their daily activities become easier. Our work on OME-Zarr to date shows an example of how community consensus and investment can achieve concrete progress towards FAIR data formats and tools.

The initial work to support OME-Zarr focused on plugins for the primary desktop visualization and analysis platforms – napari and Fiji, as well as a web browser viewer. Each new specification was implemented in these applications in order to prevent bias towards a single platform. This was the state of the ecosystem for the initial release at the end of 2021: functional with substantial language support, but insufficient adoption to consider the format mature.

In the intervening year, the number of released tools that work with OME-Zarr has increased significantly and the amount of data available is growing similarly. This trend is also visible in domains outside of bioimaging with institutes like NASA preparing for releases of their data in Zarr-based formats as part of their “Year of Open Science”^2^ (Durbin et al. 2020), (Ramachandran et al. 2021). The NGFF community finds itself in a very exciting phase. There is now a cloud-optimized, chunked format that functions as a common API for both desktop, cluster, and web-based tools as well as national and international repositories. Institutes and repositories are working towards publishing their data in a common format. For users, this means that many of their most common scalability issues can be addressed by a solution that currently exists.

At the highest level, an OME-Zarr is a single bioimaging container for multiple image acquisitions as well as derived data. The versatility of the format stems in part from the underlying data format, Zarr, and in part from the OME-NGFF community-defined specifications that are encoded in the metadata of the Zarr format, enabling use-cases across bioimaging domains. The development of Zarr features and new specifications is accelerating, but already they provide the features necessary to remove roadblocks to daily work.

### “Big Data”

OME-Zarr has been designed for performant reading and writing of large image data. This begins by storing the arrays of data in individual N-dimensional chunks. Since pixels that are shown together in viewers are stored together, they can be loaded more quickly. In a lightsheet dataset, for example, a 3-dimension region of 128×128×64 pixels might be colocated in a single atomic object. The current specification^3^ supports up to 5 dimensional images (time point, channel, z, y, x). In the forthcoming 0.5 specification, this constraint will be relaxed to allow N-dimensional arrays.

To reduce file sizes and transfer times, Zarr supports compression of each of the chunks. The compression algorithm (e.g., GZIP or Blosc (Alted 2010)) and its parameters can be configured transparently at the storage layer. The size of chunks is configurable allowing users to choose the optimal setting for a given use case to achieve a fine balancing between file size, number of files, and overall read and write speed for specific access patterns.

To allow smooth Google Maps-style zooming into large images, OME-Zarr supports storage of image chunks at multiple resolution levels. Viewers can load data at the appropriate resolution level (i.e., level of detail), which enables efficient access to data even from remote storage. Furthermore, many processing steps can be executed more efficiently on smaller representations of the data.

### Transparent organization

Another key characteristic of OME-Zarr is the ability to organize multiple such multi-dimensional pyramids into a hierarchy and attach metadata at each level of that hierarchy. This is achieved with Zarr “groups” which contain Zarr arrays and other groups in a hierarchical fashion. Metadata can be attached to each group and array separately in web-readable JSON files. These features of the Zarr format enable storing related data together, maintaining provenance information. For example, a raw image, its deconvolution, and even its segmentation can all be grouped together with the metadata defining a consistent interpretation of the data. Moreover, the community can make use of this metadata organization to flexibly store further metadata schemas. Where in OME-TIFF files, a single location is provided for storing OME-XML, OME-Zarr makes possible the storage of multiple standards such as “Recommended Metadata for Biological Images”, REMBI (Sarkans et al. 2021), “Minimum information guidelines for highly multiplexed tissue images”, MITI (Schapiro et al. 2022), or “Quality Assessment and Reproducibility for Instruments & Images in Light Microscopy”, QUAREP-LiMi (Nelson et al. 2021) alongside the OME-XML metadata.

### Collections of images

With these combined capabilities, complex, data-rich collections can be constructed to support diverse applications. The OME-NGFF specification for high-content screening (HCS)^4^, for example (Fig. 2), defines multiple levels of hierarchy for storing a plate, each of its wells and each of the fields of that well as a separate image pyramid. Similarly, segmentations, known in other domains as “annotations”, “regions-of-interest” or “segmentation maps”, can be stored as labeled images beside the raw image that was segmented. Multi-modal datasets that represent multiple acquisitions on the same sample can be stored along with location information so that the images can be overlaid on one another for visualization without changing the original underlying data.

**Fig. 2.**
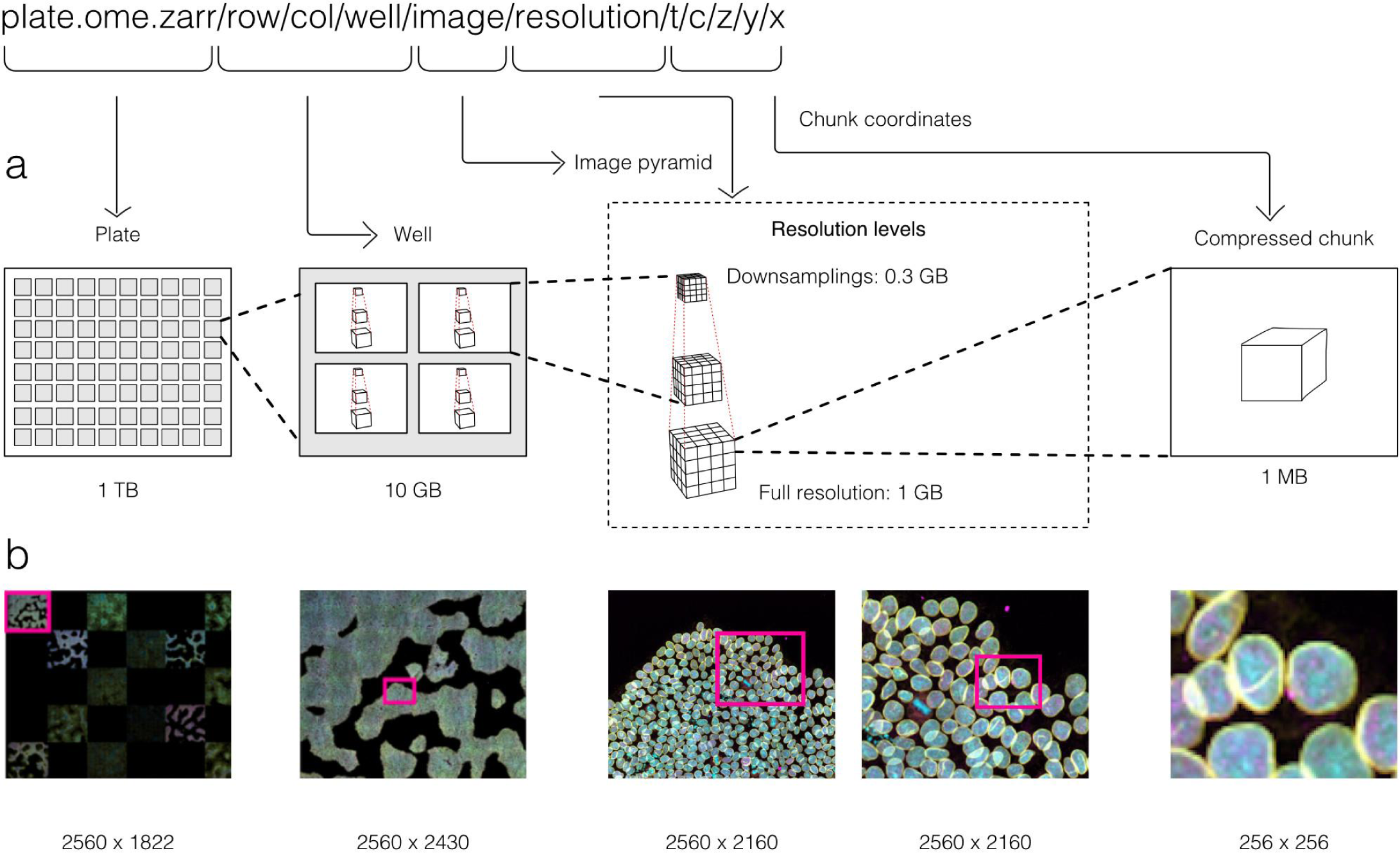
By making use of an annotated hierarchy of arrays, OME-Zarr can represent complex relationships between images, capture the multiple resolutions of an image pyramid, and provide tunable chunk size and compression all within a single abstraction layer that can be saved as a directory of files on disk or shared remotely. (A) Each level of nested directories provides a different level of abstraction: the top-level directory can represent an entire 1 Terabyte plate with more than 100,000 pixels in the X and Y dimensions, while the lowest level directory represents individual chunks of N-dimensional data as small as 1 Megabyte. (B) In the example shown, a concatenation of low-resolution images produces a 2560 pixels × 1822 pixel representation of the entire plate, followed by similar examples of how many pixels must be loaded by a client at each zoom level.

### Next steps

OME-NGFF specifications are being regularly proposed, discussed, and implemented by the community to more accurately capture bioimaging data. For example, a specification for tables that annotate label images is slated for the upcoming 0.5 version. Based on the heavily-used AnnData table representation (Virshup et al. 2021), the objective of the label table specification is to store measurements and annotations on instances in a label image, including numeric values, categorical values (e.g., strings), sparse arrays (e.g., touch matrices to encode neighborhood relationships), and dense arrays (e.g., centroid coordinates). An early prototype of this idea from HuBMAP visualizes CODEX imaging and segmentation data in combination with segmentation centroids and clustering results simultaneously with the Vitessce framework^5^ (Keller et al. 2021). Other specifications currently under consideration include more complex transformations to support correlative microscopy, efficient point cloud storage, and the versioning of data and metadata changes.

Another key next step will be how to support the NGFF model in other storage scenarios. Being based originally on the HDF5, Zarr’s compatible feature set makes the model readily transferable between the two. This would provide the user complementary approaches for balancing scalability versus complexity. On the one hand, while the internal structure of monolithic files like HDF5 are often described by complex binary data structures accessible via libraries, each Zarr chunk can be referenced via predefined, externally stable paths which provide direct access of all chunk data and metadata at each hierarchy level and can be listed by standard file browsers. With many storage backends, this strategy enables the parallel writing of large image datasets, essential for cluster and cloud-based processing. On the other hand, the potentially large number of files produced by Zarr can create problems on some file systems, generally increasing the time to list, copy, and delete data. Having support for both gives users a choice while the use of a common model in both formats increases overall interoperability.

This and future strategies for meeting user requirements will need periodic review. An upcoming version of Zarr, v3, will support a sharded layout which places a configurable number of chunks into a single shard file. This reduces the total number of files at the cost of some writing parallelism. A similar feature is available in HDF5 using “external” files and “virtual datasets” to group many separate files together. Users looking for the optimal solution will need to carefully consider the trade-offs, e.g., the impact of a multi-file format on the average consumer while existing tools are being updated.

## Selection of OME-Zarr Tools

Many common difficulties in image handling and analysis stem from both a lack of consistency and compatibility between data inputs and outputs and the resulting siloization of available tools. Without assistance, software packages are often only able to ensure compatibility with a small portion of formats. A common strategy to deal with the proliferation of file formats is to translate from one of the many current file formats on the fly. This is how open-source libraries like Bio-Formats (Linkert et al. 2010) provide access to applications as diverse as Fiji and OMERO. Translation can contribute significantly to the scalability challenge. Additionally, metadata can get lost during image translation due to opaque file structures, leaving users to provide most metadata when sharing or submitting to public resources. Sharing and re-use is complicated by disconnected images. Minimizing the number of file formats and standardizing the included metadata, in turn, fosters collaboration and stability.

The original release and publication of the OME-Zarr format was accompanied by three tools – one in Java, one in Python, and one in the web browser – that could be used to visualize and work with OME-Zarr data (Fig. 3). Over the course of the subsequent year, the number of tools has grown significantly covering additional use cases. Several of these applications originally developed their own custom format internally in order to achieve the performance they needed but have now added support for OME-Zarr allowing them to interoperate with one another.

**Fig. 3.**
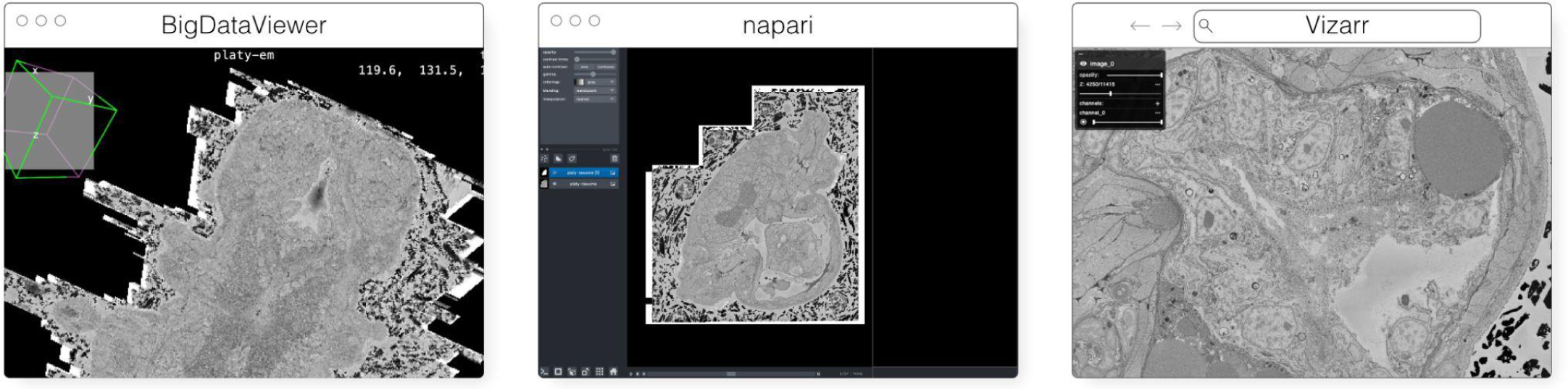
The original viewers of OME-Zarr published in Moore et al. 2021, from left to right BigDataViewer, napari, and Vizarr, here seen loading a view of the same EM volume of a 6 day old Platynereis larva from (Vergara et al. 2020) available at https://s3.embl.de/i2k-2020/platy-raw.ome.zarr. These three applications provided broad coverage over the most common bioimaging platforms like Fiji and napari but critically also a web viewer that could stream data on the fly.

Below we provide an updated list of tools that were known to handle OME-Zarr at the time of writing. This list, however, will quickly age post-publication. In order to keep track of the software packages which have added support for OME-Zarr, a registry has been created at https://ngff.openmicroscopy.org/tools. Our list is categorized into three large, though at times overlapping, categories. We start with the visualization tools (Table 1) that are broadly useful for interactively working with data. They provide an overview of what is possible with OME-Zarr. Where possible links to example data have been provided. A list of libraries follows (Table 2) that can be used to automate operations on OME-Zarr. These are useful especially when building pipelines or automating a workflow. Finally, generators (Table 4) are used to take data either from other tools or from the original acquisition system and create OME-Zarr data.

**Table 1.** List of visualization tools in the order they are described below along with their primary platform of use and the software frameworks used to build them. An up-to-date version of the table is maintained at https://ngff.openmicroscopy.org/tools and contributions are welcome.

| Visualization Tool | Use | Language/Framework |
| --- | --- | --- |
| AGAVE | Linux, MacOS, Windows | C++, OpenGL |
| ITKWidgets | Web (Jupyter) | Python, WASM |
| MoBIE/BigDataViewer | Linux, MacOS, Windows | Java |
| napari | Desktop | Python |
| Neuroglancer | Web | WebGL |
| Validator | Web | Svelte |
| Viv | Web | React, deck.gl |
| webKnossos | Web | React, WebGL |
| website-3d-cell-viewer | Web | React, TypeScript, WebGL |

### Visualization

#### AGAVE

AGAVE^6^ (Fig. 4) is an open-source native application for high quality GPU rendering of multichannel volume data. It uses a photorealistic path tracing algorithm to produce images with lighting and shadows, allowing greater detail and higher interpretability of spatial relationships within the data. AGAVE is implemented in C++ with OpenGL and runs on Windows, MacOS and Linux. OME-Zarr support is implemented through the TensorStore library, described below. AGAVE provides a memory estimate and allows selection of the multiresolution level and slice ranges in the XYZ dimensions. Future work in AGAVE will include the ability to combine OME-Zarr data from multiple data sources and improvements for more quantitative viewing such as display of physical units, voxel intensities, and a 2D slice display mode.

**Fig. 4.**
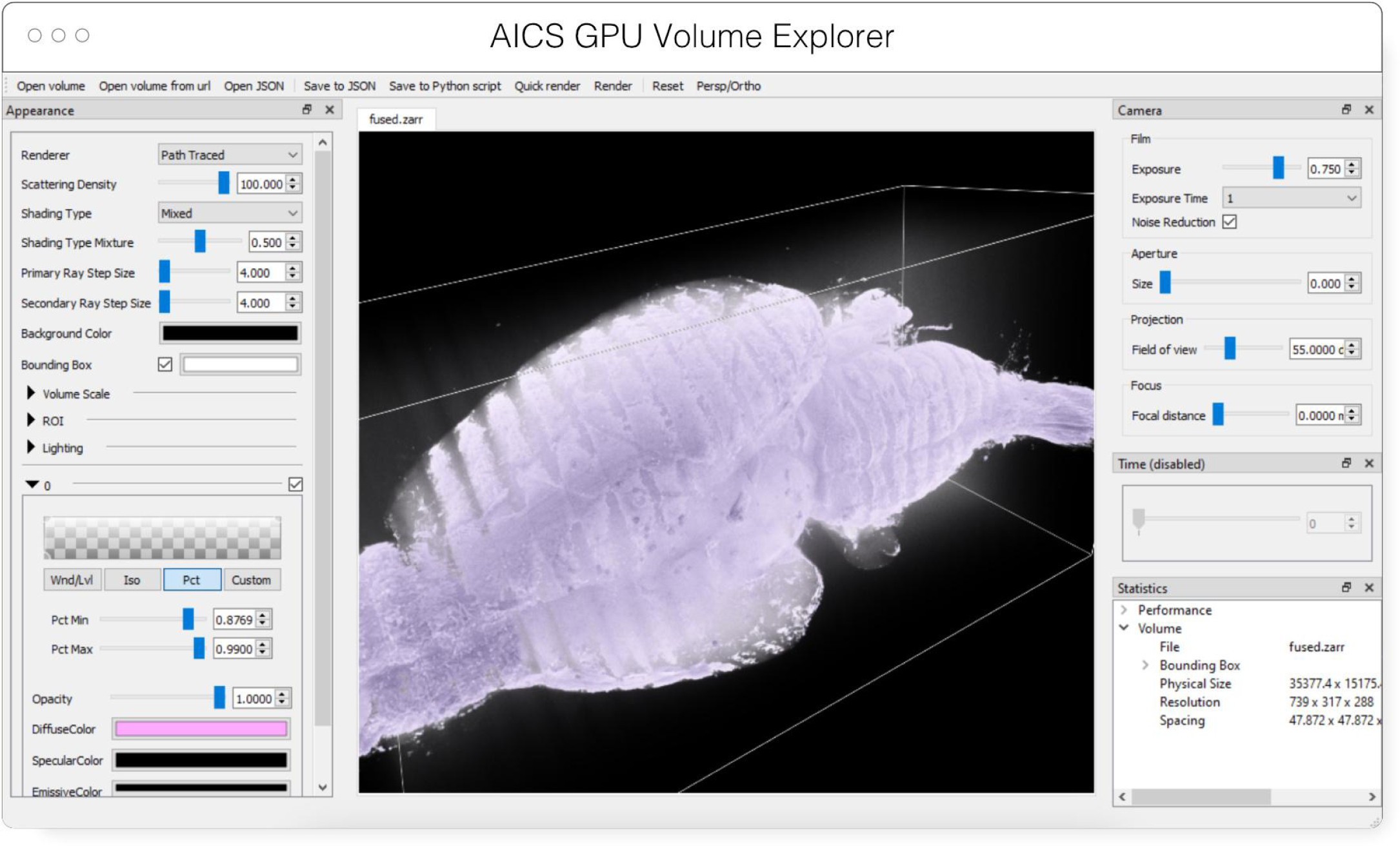
Advanced GPU Accelerated Volume Explorer (AGAVE) displaying a downsampled level from a multi-terabyte mouse brain OME-Zarr dataset. The number of pixels actually loaded is displayed at lower right. The full resolution data is 47310 × 20344 × 18471 which consumes about 33TB. The ability to quickly access multiresolution data makes low latency interactive visualization possible.

#### ITKWidgets

ITKWidgets (Fig. 5) provides interactive widgets to visualize images, point sets, and 3D geometry on the web (McCormick et al. 2022). The development of ITKWidgets was motivated by the need for interactive insights into N-dimensional scientific datasets, such as three-dimensional, multi-channel bioimages. ITKWidgets is a component of the Insight Toolkit (ITK), an open-source, cross-platform suite of libraries and tools for N-dimensional spatial analysis and visualization (McCormick et al. 2014).

**Fig. 5.**
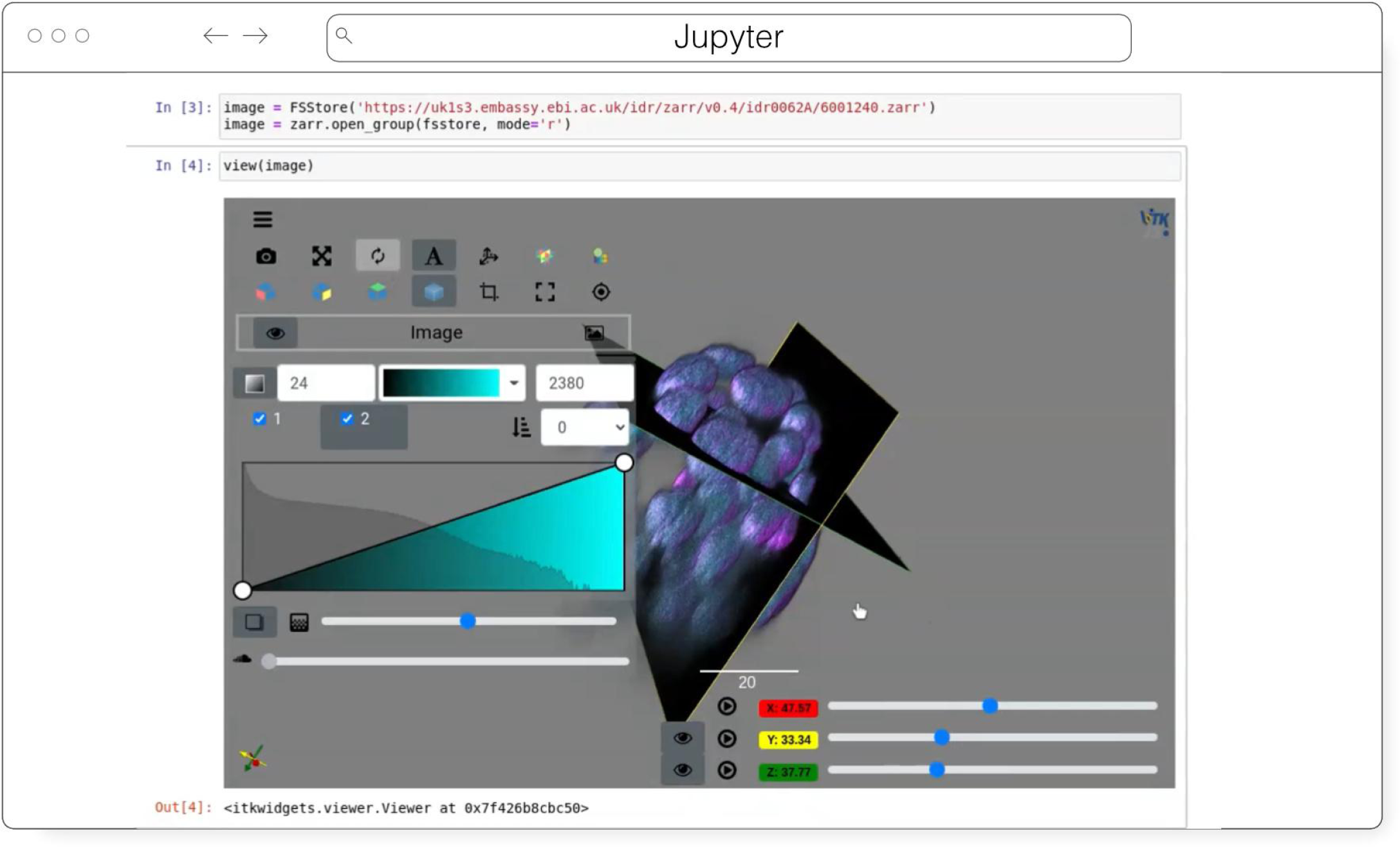
ITKWidgets 3D rendering an OME-Zarr for IDR 0062A in Jupyter. Interactive features shown include volume rendering, slicing planes, and interactive widgets to adjust rendering parameters and slice planes indices.

Designed for web-first visualization and large-scale data, ITKWidgets is built on universally deployable technologies and the OME-NGFF and ITK data models. ITKWidgets communicates with Google CoLab, Jupyter Notebooks, JupyterLab, and JupyterLite with ImJoy, a modern remote procedure communication library and plugin interface for biomedical computing in the deep learning era (Ouyang et al. 2019). ITKWidgets is available as a Python package or the client-side viewer application can be loaded by visiting its webpage. In Python, NumPy, PyImageJ, PyTorch, the Visualization Toolkit (VTK), ITK, Xarray, Zarr, and Dask data structures are transformed on-demand to multiscale OME-Zarr. In the browser, ITKWasm will generate a multiscale OME-Zarr on-demand from common bioimaging file formats (McCormick 2022). In Python, a simple *view* command accepts datasets for interactive exploration. In the client-side application, a local dataset can be selected or a shareable URL with the dataset and rendering parameters can be generated.

With a focus on supporting registration (alignment), ITKWidgets is recommended for the comparison of datasets. Spatial metadata on multi-dimensional raster images along with associated point-based volumetric data, geometries, and annotations are supported to understand their relationship in space. Additionally, this provides a foundation for the creation of spatial transformations that define or improve on the alignment of datasets. ITKWidgets is particularly focused on providing elegant renderings to elucidate insights from volumetric information. Advanced rendering capabilities, such as light scattering, are supported. Intuitive and efficient interactive widgets are available to select rendering parameters, such as color maps and opacity transfer functions. The rendering system leverages OME-Zarr chunked, multiscale architecture to automatically load the optimal amount of data for a selected volumetric region by accounting for the current system’s hardware capabilities.

The user interface is customizable via vanilla HTML/CSS/JavaScript or web frameworks such as React.js or Vue.js, and the ability to present simplified versions of current interfaces and transparently integrate the viewer into larger applications is improving. This flexibility enables integrations into custom applications such as TensorBoardPlugin3D, a plugin for TensorBoard to support the development of deep learning models on 3D images (Major and McCormick 2022). Scalability will be achieved through bolstered OME-Zarr data model support.

#### MoBIE/BigDataViewer

MoBIE^7^ (Fig. 3) is a Fiji plugin (Schindelin et al. 2012) for the exploration of big, possibly remote, image datasets (Pape et al. 2022). The development of MoBIE was initiated in 2018 at EMBL Heidelberg in order to solve the challenge of browsing and publicly sharing a large CLEM dataset consisting of one high-resolution TB sized 3D volume EM dataset, cell and tissue segmentations of the EM data, tables with segmentation annotations, and around 200 registered lower resolution LM images (Vergara et al. 2020) and is still in daily use across the institute.

The main usage of MoBIE is to continuously browse complex image datasets from the moment they are produced up until publication. A typical workflow is to use other applications for image and data analysis and add the output of those applications such as segmentations and tables into the corresponding MoBIE project for visual inspection, exploration and quality control. An exception is the possibility to perform semi-manual image registration directly within MoBIE by means of an integration with the BigWarp Fiji plugin (Bogovic et al. 2016).

MoBIE is a desktop application written in Java that heavily relies on BigDataViewer^8^ (Pietzsch et al. 2015) for image rendering and the N5 library^9^ for (remote) image I/O, described below. It supports viewing locally (e.g. file-system) and remotely (e.g. “Simple Storage Service”, or S3) hosted OME-Zarr image data as well as HDF5 and the eponymous N5 multi-scale image data format. In addition to simply viewing OME-Zarr images in Fiji, the main usage and feature of MoBIE is the ability to structure potentially thousands of images into a “MoBIE project” and define and configure useful views into that dataset. An important application of those features are “MoBIE views” that can be configured to conveniently browse the raw data associated with figures in publications.

In the future, MoBIE will support interactive deep-learning based image segmentation by means of an integration with the BioImage Model Zoo (Ouyang et al. 2022). It will also be shipped as a conda package for opening images, segmentations and tables from the command line. This will support the visual inspection of the output of image segmentation and feature extraction algorithms. Another planned feature is the rendering of the HCS specification of OME-Zarr as a plate layout.

#### napari

napari^10^ (Fig. 3) is a multi-dimensional data viewer written in Python (Sofroniew et al. 2022). Many different types of data can be viewed and annotated in napari including multi-dimensional images, point clouds, polygons, tracks, and meshes. napari can be scripted using its Python API, used interactively via interactive environments such as IPython (Perez and Granger 2007) and Jupyter notebooks (Granger and Pérez 2021), and launched from the command line. While the core napari package is focused on interactively viewing and annotating data, it can be extended to other use cases via custom scripts or through the plugin interface.

OME-Zarr data can be viewed in napari via the napari-ome-zarr plugin^11^. Users can load OME-Zarr datasets through the command line interface or via the Python API. Datasets can be loaded from both local and remote data sources. Local OME-Zarr files can also be loaded via drag & drop. Developers can use the ome-zarr-py library to load datasets and add them to the viewer via the Python API. The Fractal framework uses Dask lazy loading with the napari-ome-zarr plugin and the experimental napari asynchronous loading feature (under development, NAP-4^12^) to interactively view 2D multichannel datasets from 100s of GBs to 1TB in size (see Fractal section below). The SpatialData framework also combines Dask lazy loading and the napari plugin napari-spatialdata^13^ to visualize spatial omics data, that often entails a variety of data types: raster images, points, polygons and annotations.

#### Neuroglancer

Neuroglancer^14^ (Fig. 6) is an open-source web-based visualization tool for multi-dimensional volumetric data. Originally designed for visualizing petabyte-scale volume electron microscopy datasets of brain ultrastructure, it is now widely used to visualize scientific imaging data in many different application areas, including connectomics, lightsheet functional calcium neuroimaging, fMRI, and high-throughput screening. Key functionality includes:

- Scalability to petabyte and larger datasets through the use of multi-resolution data streaming for OME-Zarr and other chunked formats
- Cross-section views at arbitrary oblique angles
- Rendering of segmentations and meshes
- Arbitrarily many datasets may be displayed tiled side-by-side, or overlaid as separate “layers”
- Mapping from stored data to displayed RGBA values may be customized through user-defined “shader” functions that can be edited directly within the tool, and these shaders can make use of user-defined UI controls such as sliders and checkboxes
- Experimental volume rendering support
- Supports Zarr data served from arbitrary HTTP servers, as well as Google Cloud Storage (GCS) and Amazon S3

Neuroglancer is built using WebGL and relies on advanced GPU rendering and compression techniques to achieve high performance despite the limitations of the web platform. As a web-based tool, Neuroglancer is particularly convenient for collaborating on datasets; users can share particular views of a dataset simply by sharing a URL. As a purely client-side web application, Neuroglancer can be used directly from the official hosted instance, or it can be deployed to any static file web server. There is also a Python package (“neuroglancer” on PyPI) that allows for full interaction with the viewer state from Python, defining of custom key and mouse bindings that invoke Python callbacks, and also allows Neuroglancer to display in-memory NumPy arrays, as well as arrays from other packages such as TensorStore, zarr-python, Dask and h5py that provide a similar NumPy-like interface. The Python package can be used both in standalone Python programs and shells and also from Jupyter notebooks, and provides a convenient way to quickly build ad-hoc data analysis and proofreading tools.

**Fig. 6.**
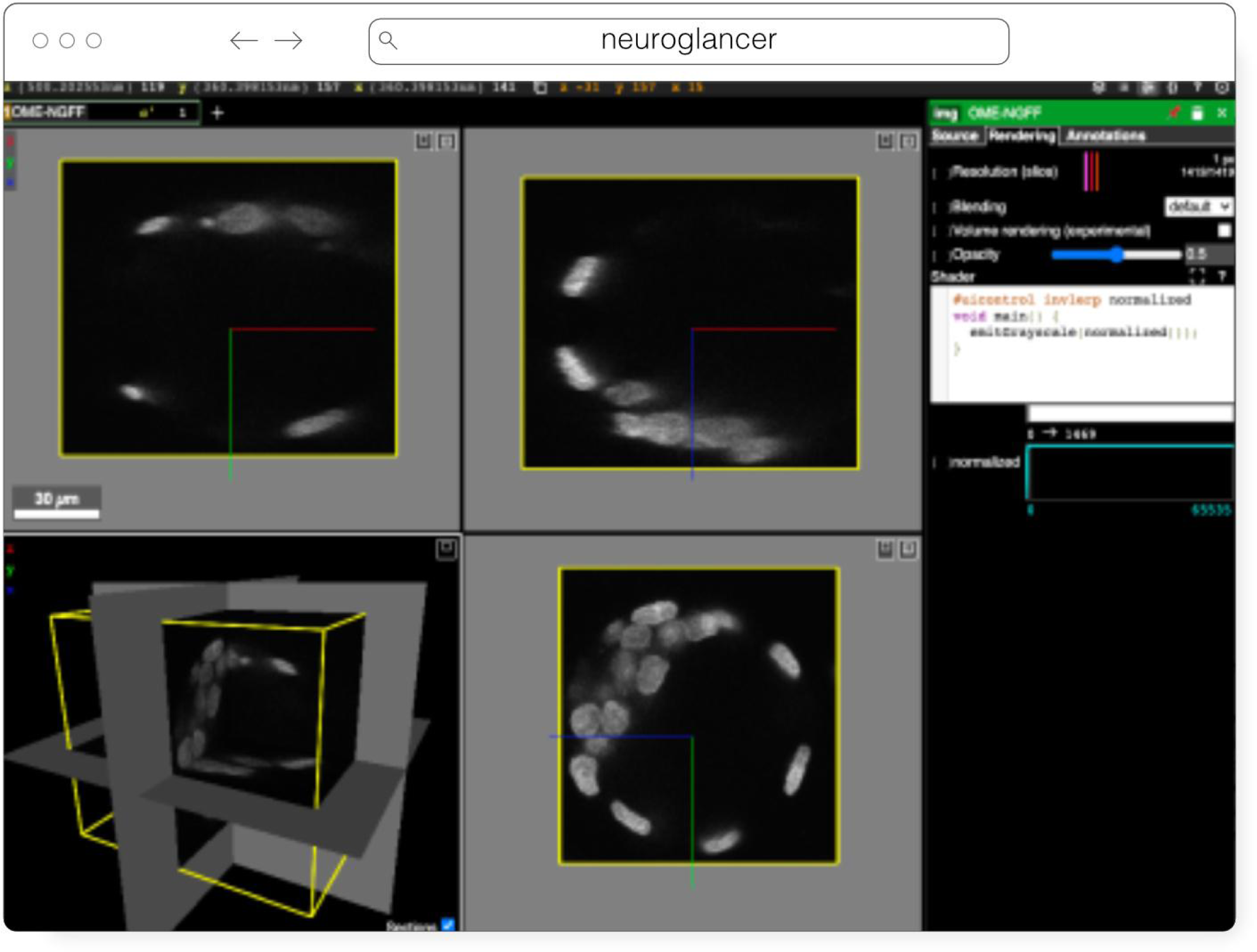
Neuroglancer rendering the same OME-Zarr from IDR 0062A as Fig. 5

#### Validator

The ome-ngff-validator^15^ (Fig. 7) is a web-based tool for validating and inspecting OME-Zarr data. It uses schema files (in the JSON schema format) for validating the JSON data of OME-Zarr, and uses zarr.js for loading image data chunks. Providing the community with an easy way to validate their data is an important part of promoting the adoption of OME-Zarr. When newly developed tools are exchanging data in this format, it is essential to know whether the data complies with the OME-Zarr specification. It is also useful to be able to browse and inspect the data in order to troubleshoot any issues with creation or reading of the format.

**Fig. 7.**
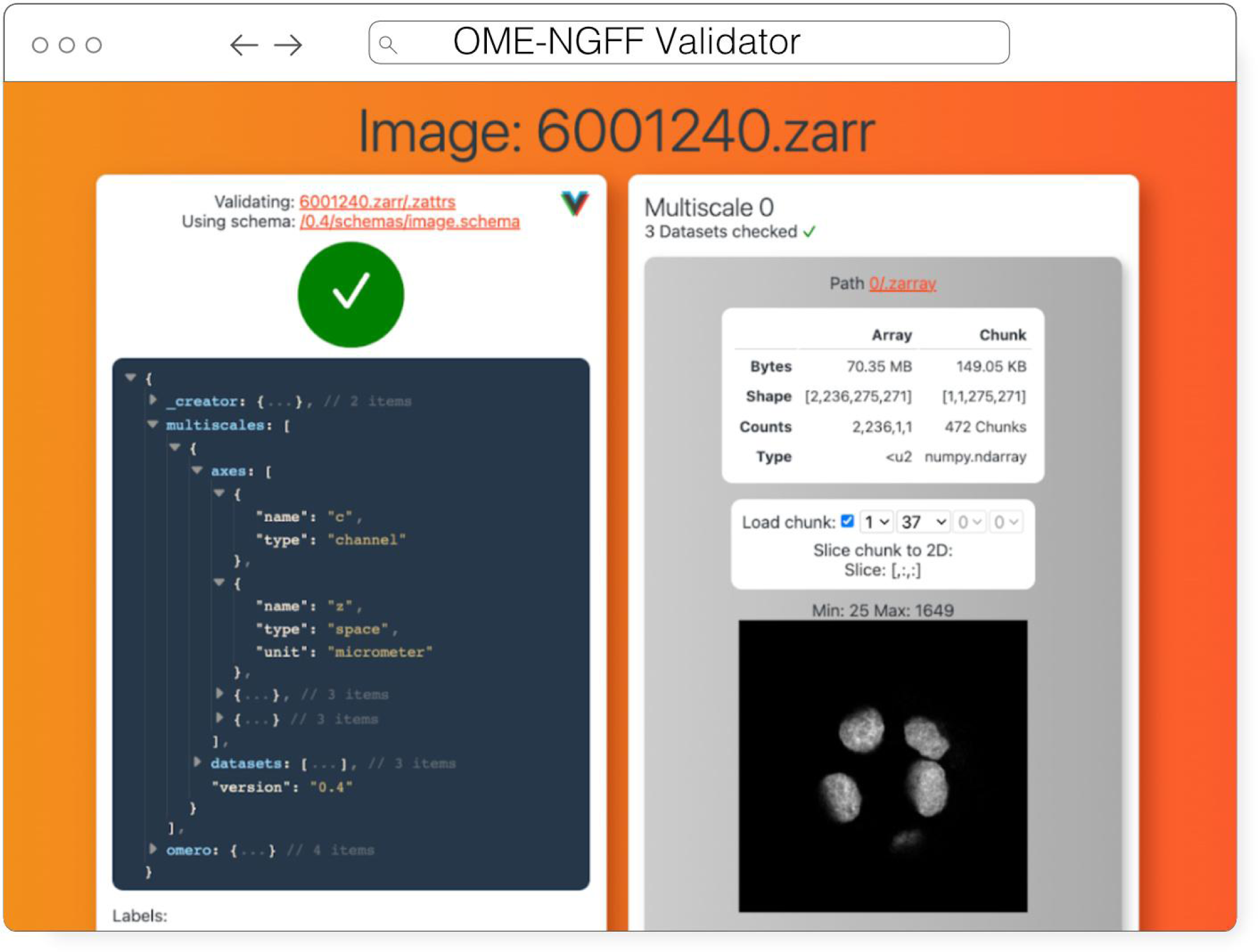
OME-NGFF Validator validating the same image from IDR 0062A on the left, and on the right providing a summary of the size of the data as well as providing a quick visualization of a single plane.

A web-based tool is convenient for users as they do not need to install any software, and it also means that they can share a URL that shows their data in the ome-ngff-validator^16^. This improves the ability of the community to discuss and work with public OME-Zarr files.

#### Viv

Viv^17^ is an open-source library for WebGL-based 2D multiscale, multi-channel, image rendering with the ability to handle 3D fixed-resolution volumes as well (Manz et al. 2022). Separation of rendering and data-loading implementations in Viv allows the library to provide functionality for fetching data directly from multiple open-standard data formats. OME-TIFF and OME-Zarr images can be loaded using Viv and directly rendered on the GPU. Viv is built using deck.gl, a high-level API for WebGL that simplifies building interactive spatial (i.e., Cartesian coordinate-based) and geospatial data visualizations. In deck.gl parlance, Viv simply provides “layers” that can be composed with other layers or operations to create complex visualizations. As a result, developers using Viv can take advantage of the wider ecosystem around deck.gl that includes custom layers, modes of interactivity, mathematical operations, and shaders. This flexibility was core to the design of the Viv API, as end-users can extend the provided layers or define custom data loaders Viv was initially motivated by the need of members of the HuBMAP consortium to display high resolution, multi-modal (i.e., overlaid) images within the HuBMAP data portal (e.g., http://vitessce.io/#?dataset=neumann-2020). Working within the constraints of limited engineering resources, the data portal development team aimed to avoid running and maintaining server-side pre-rendering infrastructure. Further, FAIR data access principles were paramount to the creation of the data portal. The development of Viv enabled rendering images within a web page, thereby enabling the data portal server-side infrastructure to be as simple as a static file server or a cloud object storage system. Adoption of FAIR data access principles in the consortium motivated the implementation of data loaders for open-standard formats. Viv has been adopted by other consortia including the Kidney Precision Medicine Project (KPMP) (de Boer et al. 2021) as well as the Galaxy Project (Galaxy Community 2022) to address data sharing and visualization challenges. Wellcome Sanger Institute in collaboration with Newcastle University is working on a human whole embryo project that leverages different spatial technologies to create a holistic view of human embryo at the single cell level. Vitessce, which is built on top of Viv, is used for visualizing both the single cell sequencing and imaging data simultaneously that are saved as OME-Zarr.

In addition to the bioimaging rendering challenges that Viv addresses in production, it has also served as a testbed and mechanism to prototype new data standards. Vizarr^18^ (Fig. 3) is a bioimaging visualization tool built using Viv that served as one of the first implementations of a reader and renderer for the HCS metadata standard in OME-Zarr^19^. Using Viv as the core rendering library, Vizarr is able to simultaneously render hundreds of images from an HCS dataset. The design of Viv as a UI-agnostic library that separates rendering from data loading means that it will remain possible to quickly develop or adapt existing applications to the evolving and increasingly flexible OME-NGFF specification.

#### webKnossos

webKnossos^20^ (Fig. 8) is a web-based, open-source platform to visualize, share and annotate very large image datasets collaboratively (Boergens et al. 2017). webKnossos was originally developed to distribute 3D image datasets of a few hundred gigabytes to students for annotation. It features segmentation tools to generate label masks for manual analysis or as training data for deep learning-based analysis. Combined with mesh rendering features, webKnossos can visualize large-scale segmentations. Additionally, webKnossos features efficient skeleton annotation tools for creating neuron traces. In many labs, webKnossos has become the primary hub to store all image datasets in a common place for lab members and external collaborators to access.

**Fig. 8.**
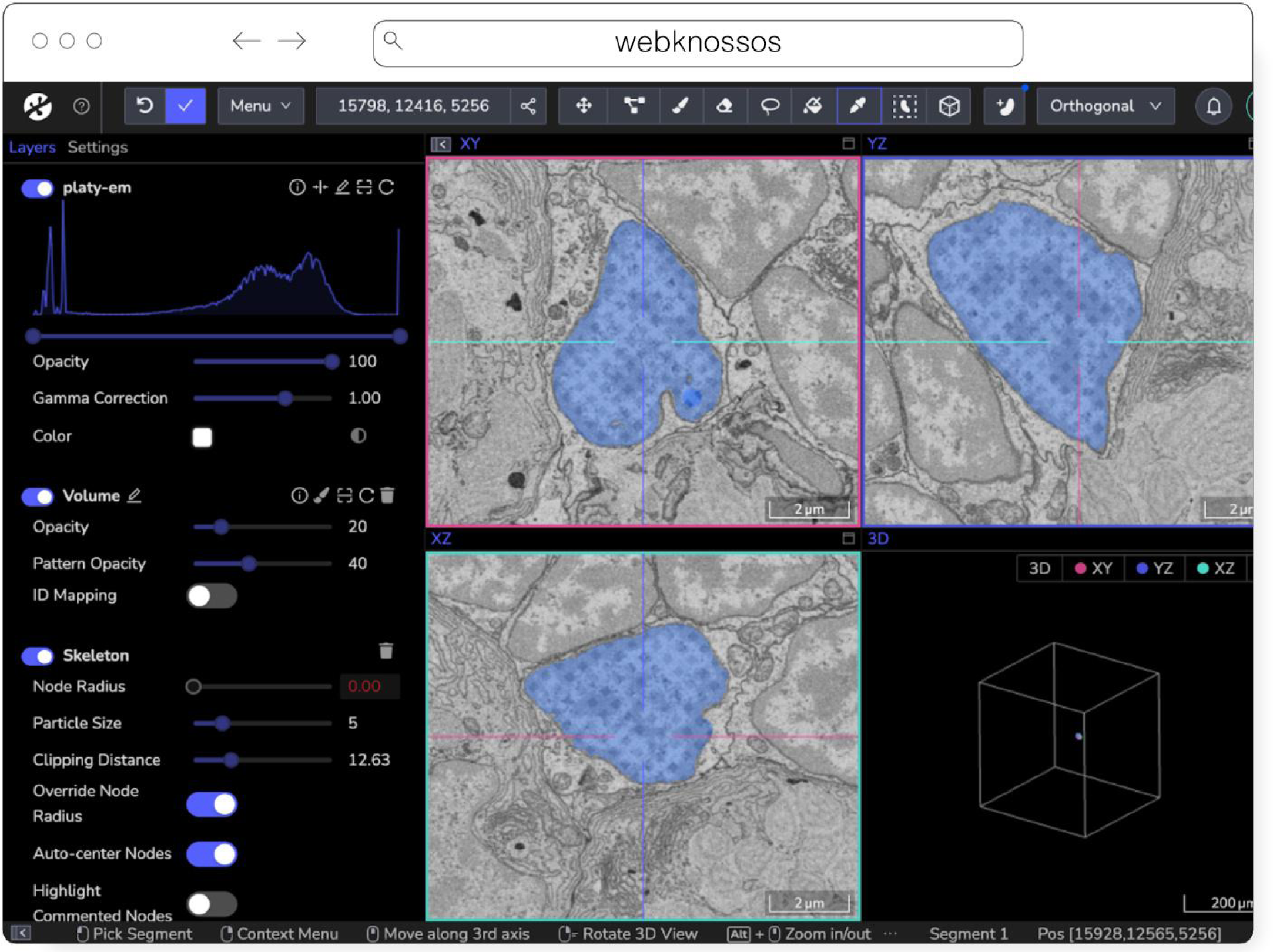
webKnossos loading an EM volume of a 6 day old Platynereis larva from (Vergara et al. 2020) with a manually added segmentation. The web accessible version is accessible at https://wklink.org/6422.

webKnossos works best for very large 3-dimensional image data, such as EM, MicroCT and fluorescence. It facilitates collaboratively annotating and visualizing data. Users can:

- visualize very large 3D datasets,
- manually segment or skeleton data efficiently,
- proof-read automatic segmentation,
- share data with collaborators,
- manage data in one place with access restrictions, and
- stream data through an OME-Zarr-based API or use a Python library to enable interoperation with other tools.

webKnossos allows users to store all data in one place and access it from wherever you are, no matter the size of the data, like a Google Drive for large microscopy images. The server component stores all the image and annotation data as well as user settings. In addition to server-stored image data, remote datasets stored in OME-Zarr can also be accessed from HTTP, S3 and GCS sources. webKnossos uses GPU rendering for high-performance visualization of the data. Users can access webKnossos through a browser without the need for additional installations. The easiest way to use webKnossos is to open an account on webknossos.org. It is free for limited storage. Alternatively, both a Docker-based setup and commercial private hosting services are available.

webKnossos is a mature software and in routine use for more than 10 years. It gained support for OME-Zarr in 2022 and implements many features of the format. OME-Zarr datasets can be imported via URL, optionally with credentials. OME-Zarr data is read on the backend with on-the-fly re-chunking and additional webKnossos-managed access controls. Additionally, all data in webKnossos is accessible as OME-Zarr data to be used in other tools via token-protected dynamic URLs. Therefore, the software can be used as a central hub for teams to manage OME-Zarr datasets. webKnossos is actively developed by a dedicated development team with monthly releases. Today, most users are from the Volume EM community, especially EM connectomics and cell biology (Rzepka et al. 2023). However, light microscopy users are also well-represented. In upcoming versions, webKnossos will add support for image transformations, multi-modal datasets, and time-series data as well as the ability to run AI-based segmentations and to show segment statistics for quantitative analysis. Updated roadmap information is available under https://webknossos.org/roadmap.

#### website-3d-cell-viewer (and volume-viewer) from AICS

The Allen Institute for Cell Science is an open science organization, interested in producing results, data, and code that can be shared with the world. Volumetric microscopy datasets are presented for interactive exploration through the Cell Feature Explorer^21^, a web-based application. As with so many microscopy efforts, ever larger and larger datasets are the norm. As an early part of transitioning storage and processing to the cloud, the interactive 3D viewer component of Cell Feature Explorer has been extended to read OME-Zarr data.

The core of the viewer component is called volume-viewer which implements all of the data loading and 3D rendering, using WebGL and Typescript (Fig. 9). The front end, called website-3d-cell-viewer, is a reusable React component that has also been published as a standalone viewer^22^ and embedded in a Jupyter notebook plugin^23^.

**Fig. 9.**
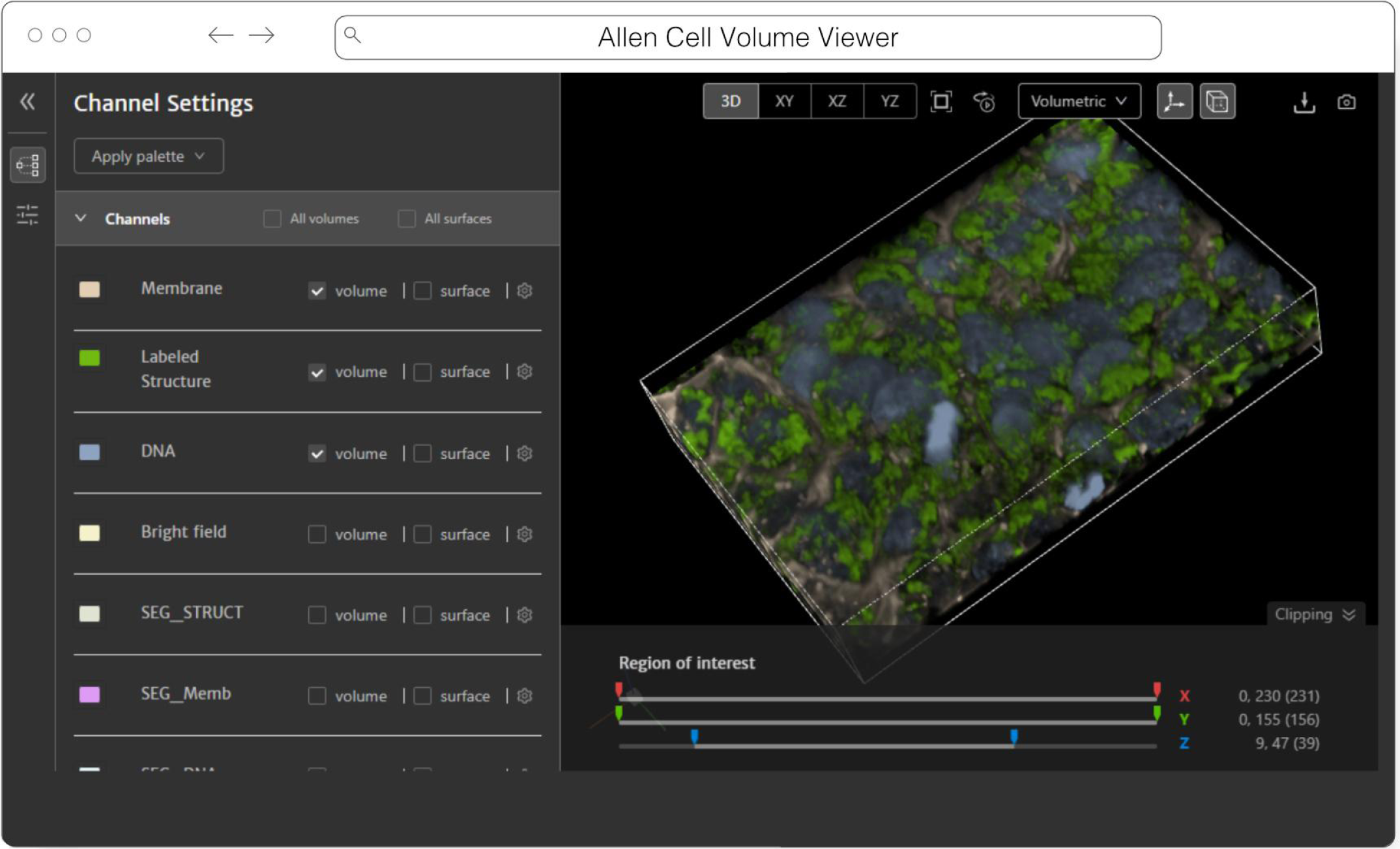
Allen Cell Volume Viewer displaying a multichannel fluorescence image of gene edited hiPSC cells via a downsampled level of an OME-Zarr converted from the dataset found at https://cfe.allencell.org/?dataset=variance_v1

This viewer is optimized for multichannel data and works well with volumetric data where the xy resolution is higher than z. The standalone version of the viewer supports OME-Zarr through a URL query parameter. The OME-Zarr support is implemented using the Zarr.js library^24^. In its first implementation, only TCZYX data is supported, only from publicly accessible HTTP(S) URLs, and as of this writing only loads the lowest multiresolution level for memory constraints. Coming enhancements include more general data loading as well as user interface for selection of multiresolution level and time. There is also work being done to be able to produce shareable URLs that capture all viewer settings.

#### Beyond

More analysis tools leveraging the libraries to create derived datasets and produce quantitative insights into OME-Zarr data are already planned. For example, the BigStitcher (Hörl et al. 2019) toolchain which currently relies on a custom XML/HDF5 format will gain support for OME-Zarr, making it easier to parallelize distributed access for HPC and avoid additional file conversion. This is particularly important for very large datasets and means that NGFF will be natively exported by BDV in Fiji. Work on this is underway in the bigdataviewer-n5^25^ and bigstitcher-spark^26^ repositories. At the time of writing, BigStitcher has support for large file export and limited direct support of OME-Zarr datasets via the bigdataviewer-omezarr module^27^. This module provides a transient bridge between the current XML-based dataset definition and OME-Zarr images until native support is implemented. It allows multiple images stored in one NGFF dataset to be defined as separate view setups in BDV/BigStitcher (Fig. 10). Other tools such as RS-FISH (Bahry et al. 2022) for spot detection and STIM for visualizing and reconstruction of spatial transcriptomics datasets (Preibisch et al. 2022) are currently being extended to support OME-Zarr and other formats. Additionally, registration tools that generate the developing spatial transformation specification are planned to enable quantitative comparison of compatible datasets.

**Fig. 10.**
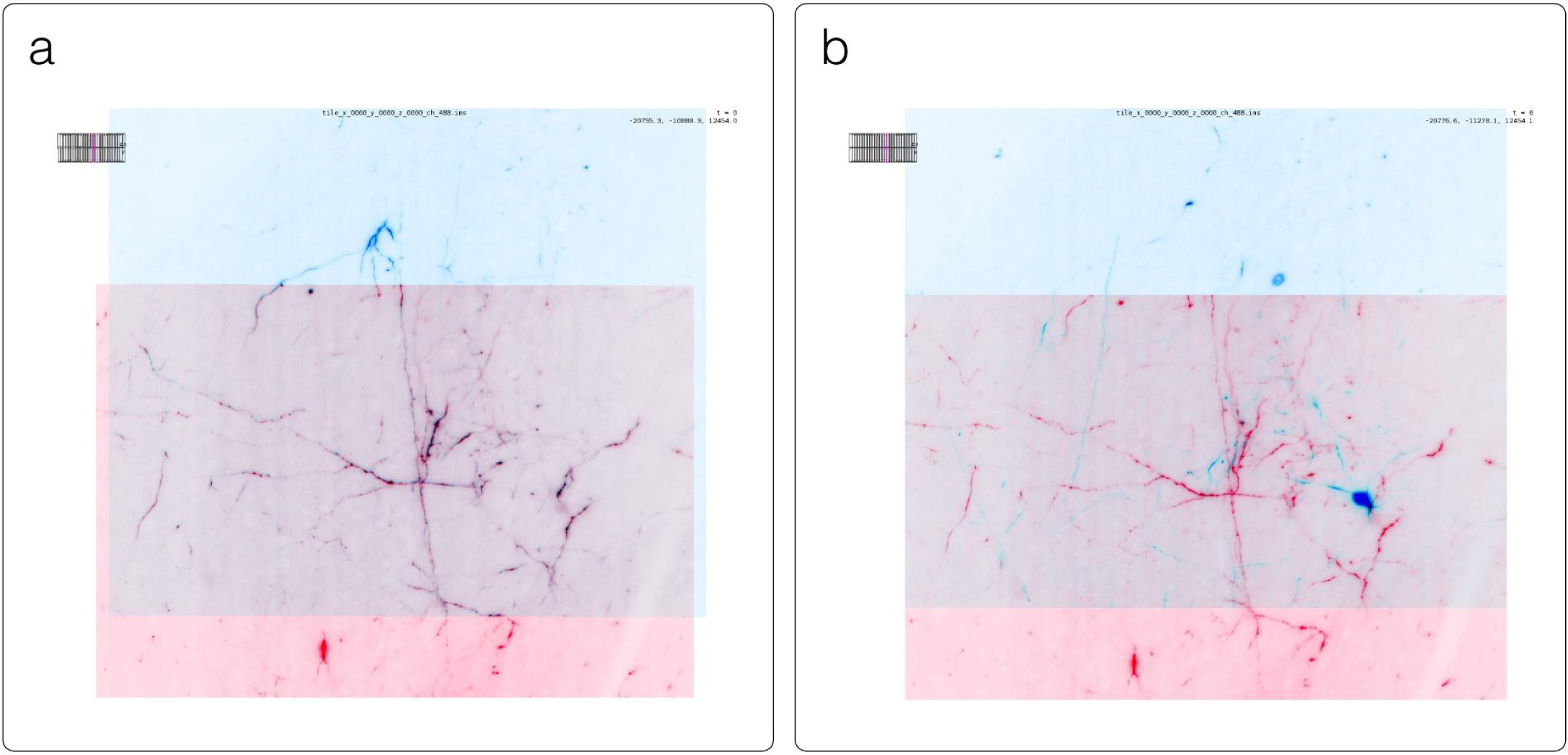
Example application of BigStitcher’s interest point based registration on one of the first “exaSPIM” lightsheet microscope datasets acquired at the Allen Institute for Neural Dynamics (sample 609281, available at the link in Table 2). The overlapping regions of two tiles are shown in BDV at their nominal (left) and aligned (right) locations. This large scale NGFF dataset consists of 54 tiles with dimensions of 24576 × 10656 × 2048 voxels (about 1 TB raw size) each.

To track these developments, the community will maintain https://ngff.openmicroscopy.org/tools for finding the status of other new and exciting developments.

#### Libraries

Behind most of the visualization tools above and many other applications are OME-Zarr capable libraries (Table 2) that can be used in a wide variety of situations. Workflow systems like Nextflow or Snakemake can use them to read or write OME-Zarr data, and the same is true of machine learning pipelines like Tensorflow and PyTorch. Where dedicated widgets like ITKWidgets are not available, these libraries can make use of existing software stacks like Dask and NumPy to visualize the data in Jupyter Notebooks or to perform parallel analysis.

**Table 2.** List of libraries broken down by programming language in the order they are described below along with a brief description of their use. An up-to-date version of the table is maintained at https://ngff.openmicroscopy.org/tools and contributions are welcome.

|  |  |  |
| --- | --- | --- |
|  | BFIO | Optimized reading and writing of TIFF and Zarr |
|  | SpatialData | Enable the alignment of spatial omics datasets |
| **C++** | TensorStore | High-performance access to multiple array formats |
|  | Nyxus | Out-of-core, parallel feature extraction library |
| **Java** | OMEZarrReader | Plugin for reading OME-Zarr in Bio-Formats |
|  | N5 API | Array data reading of Zarr, HDF5, and N5 formats. |

#### Python

ome-zarr-py^28^, available on PyPI, was the first implementation of OME-Zarr and is at the time of writing considered the reference implementation of the OME-NGFF data model. Reading, writing, and validation of all specifications are supported, without attempting to provide complete high-level functionality for analysis. Instead, several libraries have been built on top of ome-zarr-py. AICSImageIO is a popular Python library for general 5D bio data loading. In addition to loading OME-Zarr data, AICSImageIO provides OmeZarrWriter using ome-zarr-py under the hood. In this way, format conversion is possible by loading data with AICSImageIO and immediately passing the Dask array to OmeZarrWriter, though improvements in the metadata support are needed.

Fractal^29^ is a framework to process high-content imaging data at scale and prepare it for interactive visualization. Fractal is focused on processing images in the OME-Zarr format and saving results in forms of images, label images and feature tables back into OME-Zarr, while keeping orchestration of these processing steps cluster friendly via slurm^30^. It allows users to build workflows to process images in OME-Zarr files at the TB scale to segment images and make measurements. As a result, large-scale OME-Zarr image datasets can be processed by Fractal and then browsed interactively across pyramid levels in viewers like napari^31^ (Fig. 2). Fractal is in its early stages and currently contains workflows to be controlled from the command line interface. A web client to build workflows and manage the processing is currently being built.

BFIO^32^ is a Python library that supports reading of all 160+ Bio-Formats supported file formats, with opinionated but highly optimized reading and writing of OME-TIFF and OME-Zarr for use in large scale applications. All changes and updates to the specification are implemented in the library. Similar to AICSImageIO, BFIO can act as a format conversion tool where it is possible to load data in diverse image formats and immediately pass the NumPy array to a custom OME-Zarr writer function. BFIO distinguishes itself from other NGFF loaders in that it has focused on performance and performs chunked data read/writes to load and save images in an out-of-core fashion by default. BFIO is opinionated about chunk size and loading/saving pattern and this loss of freedom by the user allows BFIO to make substantial gains in its read/write speeds. BFIO is currently in the process of being refactored to utilize TensorStore with subsequent additions to TensorStore to support the full OME data model, and is planned to release the optimized TensorStore read/write in the coming months.

SpatialData^33^ is a Python library that provides an on-disk format and an in-memory object for working with spatial omics data (Fig. 11). It uses the ome-zarr-py readers and writers for raster data (images and labels) and implements features of the upcoming version of the NGFF specification, including tables and transformations. Furthermore, it experimentally supports the representation of additional modalities commonly found in spatial omics data, such as points (e.g. transcripts locations) and shapes (e.g. spatial transcriptomics circular array capture locations and generic polygonal ROIs). The library implements a set of operations such as aggregation of molecular measurements (associating transcripts locations to cells) and efficient spatial queries that are interoperable across representations. Finally the SpatialData framework provides readers for datasets produced by the most popular spatial omics technologies, a static plotting library based on matplotlib (Hunter 2007) as well as a napari plugin for interactive visualization and annotation of the data.

**Fig. 11.**
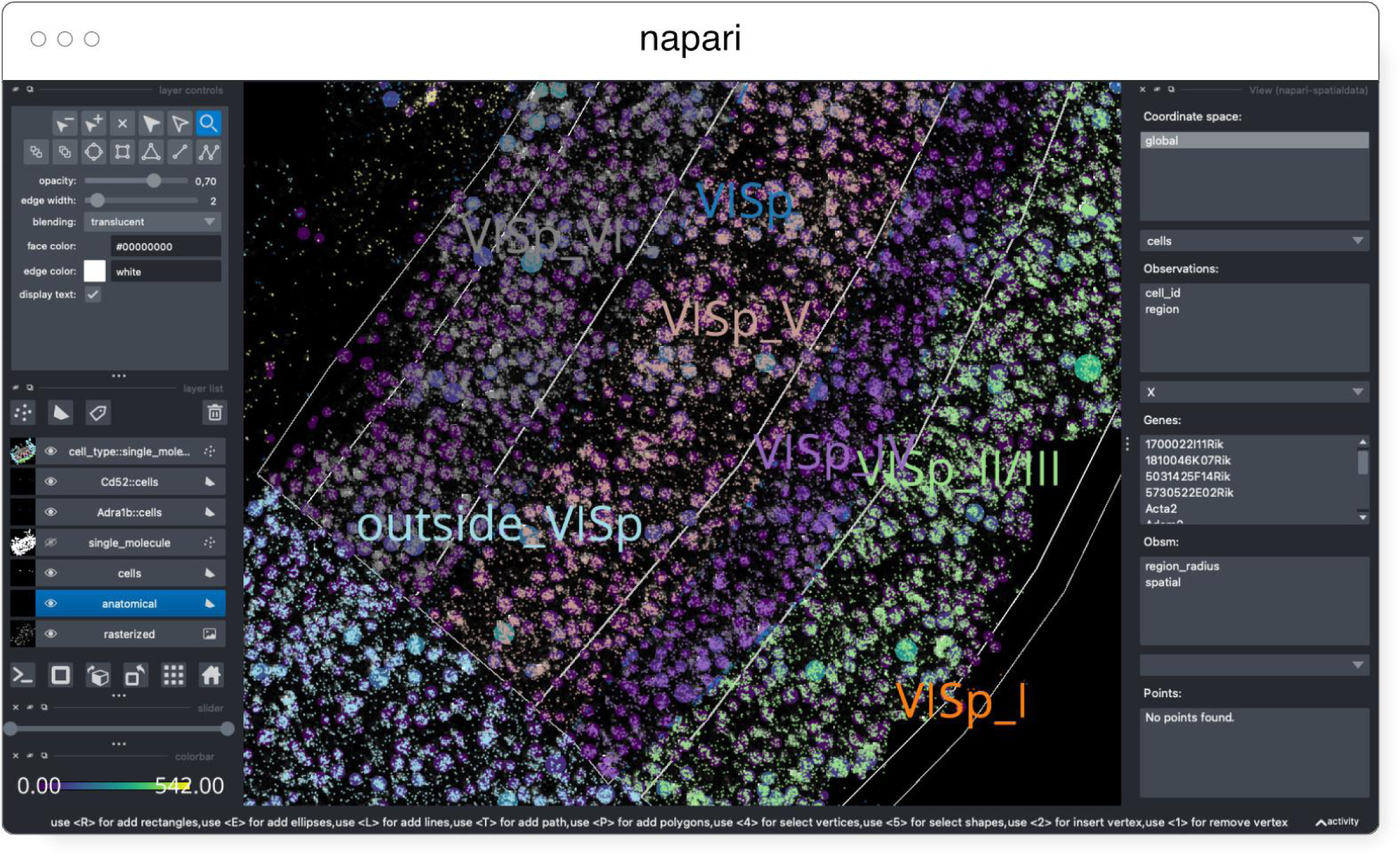
Visualization of a MERFISH mouse brain dataset (Allen Institute prototype MERFISH pipeline (Long et al. 2023)) via the napari-spatialdata plugin, featuring single-molecule transcripts (points) and their rasterized representation (image), polygonal ROIs, and annotated cells approximated as circles with variable radii. The dataset has been converted to OME-Zarr with the SpatialData APIs.

#### C++

TensorStore^34^ is an open-source C++ library and Python wrapper that provides a unified, high-performance interface for accessing a variety of array formats, including Zarr. In addition to use with scientific imaging data, it is also used to store and load checkpoints of massive machine learning models with hundreds of billions of parameters. Supported underlying storage mechanisms include: arbitrary HTTP servers (read-only); local and network filesystems; and Google Cloud Storage. It supports safe, concurrent write access from multiple machines, without the need for a separate lock server, through the use of optimistic concurrency. While it does not yet specifically support OME-Zarr multiscale metadata, it can be used to read and write individual scale levels as Zarr arrays. Specific abstractions for multi-resolution data, along with support for OME-Zarr multiscale metadata, is planned.

Nyxus^35^ is an open-source C++, plus Python wrapper available on Conda or PyPI, that provides a high performance and out-of-core, parallel feature extraction library for image and annotation data natively in OME-Zarr. The library assumes that regions of interest are labeled and can be stored as an annotation layer within OME-Zarr files or stored as separate OME-Zarr files from the raw intensity data. Nyxus supports feature extraction on 2-dimensional or 3-dimensional data and contains more morphological, histogram, and texture features than most individual libraries. It can compute these features in an out-of-core fashion enabling users to analyze images/volumes of unlimited size. Nyxus supports whole image or typical region based feature extraction, it also supports nested feature extraction (i.e. parent annotation with children annotations) and a single region across many channels.

#### Java

For reading OME-Zarr data with Bio-Formats in Java, the OMEZarrReader^36^ has been developed. Once installed, this plugin allows opening OME-Zarr images from any application that currently makes use of Bio-Formats plane-based API. This includes ImageJ or RBioFormats for accessing OME-Zarr data in R.

The N-dimensional N5 API supports reading array data and arbitrary metadata using Zarr, HDF5 and the N5 format. Its associated Fiji plugins^37, 38^ can therefore read pixel / voxel data from OME-Zarr containers, but not its metadata specification at the time of this writing. The developers have committed to adding OME-Zarr support in the future by developing a shared implementation with MoBIE. Support for multiple backends makes the N5 API an appealing choice for extending NGFF support to other, write-optimized storage formats.

#### Example

An example analysis of a dataset from the Image Data Resource (IDR) (Williams et al. 2017) helps to illustrate the possibilities available to the end-user from an analysis perspective. A light sheet fluorescence microscopy image published by McDole et al (idr0044)^39^ is composed of 2 channels, approximately 1000 z-sections and 500 timepoints. Each 2D-plane is of dimension 2k × 2k. Numerous z-sections were acquired but the relevant planes are the middle z-sections. Such data is particularly useful for teaching and training purposes, so it is usually only necessary to access a limited subset of an image.

Traditionally, two options are available in order to analyze an image. One can download the full image. This approach is far from ideal since only a portion of the data is required for analysis. Additionally, a specific image reader is needed to interpret the data, potentially limiting which analysis language/framework could be used to perform the analysis. Alternatively, the relevant planes could be retrieved using the IDR Application Programming Interface (API). The IDR API is very versatile but it complicates the parallelisation of tasks for users. To enable streamable access by all of the tools and libraries outlined above, the images of the study were converted using bioformats2raw (see the “Generators” section) and made available on object storage at EBI (See Table 3).

**Table 3.** Comparison of access methods to data stored in the IDR. OME-Zarr provides the fastest and most flexible access when accessing less than an entire dataset. A notebook documenting the OME-Zarr access is available at https://github.com/ome/omero-guide-python/blob/master/notebooks/idr0044_zarr_segmentation_parallel.ipynb.

|  |  |  |  |
| --- | --- | --- | --- |
|  |  |  | da.from_zarr(endpoint_url) |
| Easily analyze in parallel | No, depends on file format which may require a translation library. | Difficult due to the transfer protocol used (zeroc-ice) | Yes. Use Dask schedulers:<br>dask.delayed(analyze)(t, c, z) |

The nature of OME-Zarr allows the end-user to take full advantage of libraries like Dask^40^, a free and open-source parallel computing library that scales the existing Python ecosystem. An analysis task like segmentation is broken into many individual steps that are run independently. Each step lazily loads the chunk of the image that it is to work on, and then the result is aggregated. Available public resources like Google Colab^41^ or mybinder^42^ and publicly accessible data were sufficient for training purposes but the approach could easily be extended for larger scale analysis. This facilitates training the next generation of scientists on how to use cloud-computing resources.

#### Generators

For the foreseeable future, datasets will exist in one of the many forms they exist in today. As outlined in (Moore et al. 2021), translating those on the fly brings delays in visualization and analysis that can be solved by performing a single conversion to OME-Zarr. This process captures all metadata that Bio-Formats is aware of in an open format, and decouples the user from the version of the vendor software used to capture the data. Several generation tools (Table 4) are available based on the particular environment you are working in.

**Table 4.** List of software for generating OME-Zarr data in the order they are described below along with a brief description of their use. An up-to-date version of the table is maintained at https://ngff.openmicroscopy.org/tools and contributions are welcome.

| Storage | Catalog | Dashboards / Datasets | Zarr Files | Size |
| --- | --- | --- | --- | --- |
| **Amazon S3** | Cell Painting Gallery | https://github.com/broadinstitute/cellpainting-gallery | 136 | 20 TB |
|  | DANDI | https://dandiarchive.org/dandiset/000108<br>https://identifiers.org/DANDI:000108<br>https://github.com/dandisets/000108 | 3914 | 355TB |
|  | Neural Dynamics | https://registry.opendata.aws/allen-nd-open-data/ | 90 | 200TB |
|  | Glencoe | https://glencoesoftware.com/ngff | 8 | 165 GB |
| **Alternative S3 Providers** | BIA Samples | https://bit.ly/bia-ome-ngff-samples | 90 | 200GB |
|  | IDR Samples | https://idr.github.io/ome-ngff-samples/ | 88 | 3TB |

<sup>46</sup> https://github.com/fsspec/kerchunk

#### Bioformats2raw

Bioformats2raw^43^ is the original command-line utility to convert various image file formats into OME-Zarr format. Bioformats2raw offers rich and flexible parameter options, giving the user extensive control over the conversion process as well as freedom to specify various features of the output Zarr datasets. A few of the interesting input parameters are the chunk dimensions, the number of resolution levels, the compression algorithm, and the number of workers. Bioformats2raw can read all proprietary file formats supported by Bio-Formats as well as a select few file format readers supported only in bioformats2raw, including the 3D-Histech .mrxs format. The input, therefore, can be single or multiple series as well as high-content screening (HCS) data. The conversion will be performed according to the respective OME-NGFF specification. Multiscaling is achieved either by creating sub-resolutions during the conversion process, or by using the existing ones from the input format.

Bioformats2raw is optimal for remote and headless operation and can be conveniently built into pipelines, e.g., by using workflow management systems, such as Galaxy, Nextflow, Snakemake, etc. that would also facilitate parallel conversion of batches of image data into OME-Zarr, for example on HPC clusters. Additionally, cloud-aware tools like Distributed-OMEZarrCreator (Weisbart and Cimini 2022) allow easy wrapping of bioformats2raw on Amazon Web Services (AWS). By default, bioformats2raw writes an OME-XML metadata to a specific directory in the output Zarr hierarchy. This metadata can then be used by a complementary package, namely raw2ometiff, to convert the output from bioformats2raw into OME-TIFF.

#### NGFF-Converter

Glencoe Software’s NGFF-Converter^44^ (Fig. 12) is an easy-to-use and intuitive graphical user interface (GUI) application supporting conversion of any format readable by Bio-Formats, as well as the additional readers built into bioformats2raw. By packaging the command line utilities bioformats2raw and raw2ometiff, NGFF-Converter can convert numerous file formats to both OME-TIFF and OME-Zarr based on the user selection. NGFF-Converter is approachable for users less familiar with command line utilities while maintaining the flexibility of tunable parameters as described in the previous section. In addition, NGFF-Converter was developed with batch processing in mind, supporting the scheduling of multiple conversions with clear visuals of conversion job status. NGFF-Converter is available for both Windows and MacOS.

**Fig. 12.**
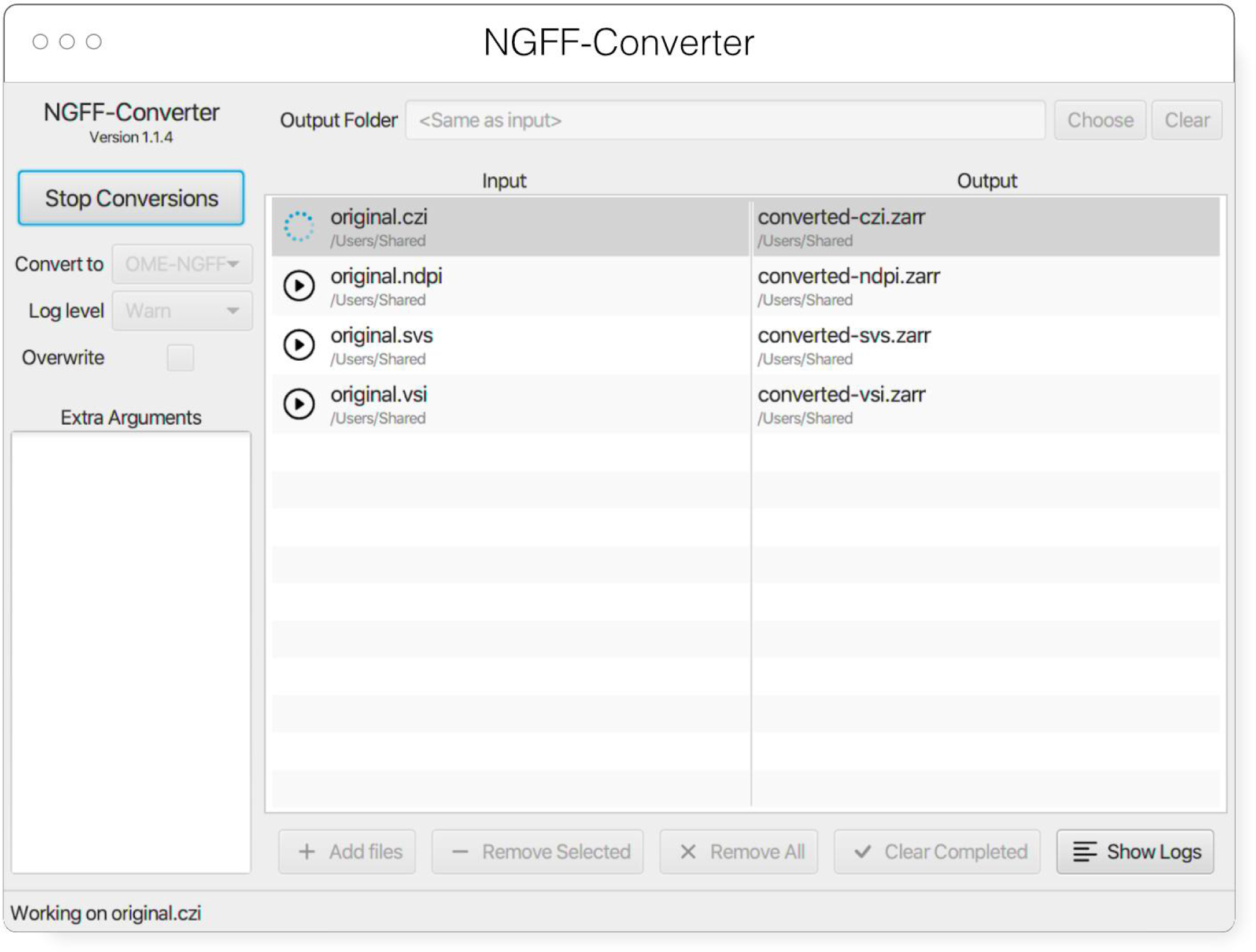
NGFF-Converter GUI showing a sample of input formats being converted to OME-Zarr

#### ImSwitch

At the same time, we envision an increasing number of hardware devices capable of directly outputting image data in OME-Zarr, streamlining analysis and reducing the risk of data duplication. One example of software that facilitates this approach is ImSwitch^45^, a modular open-source microscope control software written in Python. Imswitch implements an architecture based on the model-view-presenter design pattern to enable flexible and modular control of multiple microscope modalities. The experimental data acquired, with related experiment metadata, can be directly written with ome-zarr-py into OME-Zarr files. The file format showed to be favorable when using workflows involving distributed computational resources for processing (e.g. parallel RESOLFT reconstructions) of image data (Casas Moreno et al. 2023). Previously used acquisition parameters and settings can be loaded into ImSwitch from saved data to conveniently enable reproducible workflows.

#### Kerchunk

Within the Python ecosystem, it is also possible to “simulate” an OME-Zarr dataset with kerchunk^46^. This library pre-processes existing data formats like TIFF and HDF5 to generate a JSON file containing all metadata and chunk locations. Support for other file formats can be added if each chunk can be represented by a combination of path to a file, location in that file, and length of the chunk. Using this mechanism, it is possible to leave data in a monolithic format but still achieve some of the benefits of OME-Zarr. Support in other programming languages is possible based on community interest.

#### Other ways to create OME-Zarr

In addition to using these dedicated generators, many of the general-purpose tools mentioned also support the generation of OME-Zarr data. Within the Fiji ecosystem the MoBIE plugin provides a GUI for creating OME-Zarr. Among its other functionalities, MoBIE can convert images imported by Fiji into OME-Zarr. The input is imported via Fiji readers, which include Bio-Formats, and enables immediate visualization and exploration options of the OME-Zarr data. All uploads to webKnossos from all supported formats are also automatically converted into OME-Zarr, which can be streamed or downloaded for use with other tools. Libraries like ome-zarr-py can write numpy and Dask arrays to OME-Zarr according to the OME-NGFF specification. Where users are already manually handling the reading of the input data and the parsing of the metadata in Python code, this may be the easiest path to generating OME-Zarr data.

## Examples of Shared OME-Zarr Data

Not just individual users or research projects are faced with the issues of format compatibility. Large-scale bioimaging resources are also moving to OME-Zarr to ease access across a range of storage options. Below we discuss several of the ways in which these and other institutions are sharing their data with the OME-Zarr format as examples of what is possible as well as where you might find existing data today, all summarized in Table 5. However, as with the tools above, these and other resources are being actively updated. Users interested in re-using datasets can refer to https://ngff.openmicroscopy.org/data for an up-to-date version of this table maintained by the community or submit new resources as they become available. Though central registry does not exist for other file formats, the ease of access to OME-Zarr on the web, e.g. through embedded multiple-terabyte data in a static webpage, makes such a catalog particularly valuable and the growing availability of OME-Zarr formatted data will hopefully accelerate tool development.

**Table 5.** List OME-Zarr resources data belonging to the authors, broken down by storage types in the order they are described below along with rough estimates of their size at the time of publication. These include catalogs and dashboards which will help the reader discover datasets as well as some resources which are migrating to OME-Zarr. An up-to-date version of the table is maintained at https://ngff.openmicroscopy.org/data and contributions are welcome.

| Storage | Catalog | Dashboards / Datasets | Zarr Files | Size |
| --- | --- | --- | --- | --- |
|  | Sanger | https://www.sanger.ac.uk/project/ome-zarr/ | 10 | 1TB |
|  | SSBD | https://ssbd.riken.jp/ssbd-ome-ngff-samples | 12 | 196GB |
| **On-premise** | CZB-Zebrahub | https://zebrahub.org | 5 | 1.2 TB compressed |
|  | MoBIE | https://mobie.github.io/specs/ngff.html | 21 | 2 TB |
|  | SpatialData | https://github.com/scverse/spatialdata-notebooks/tree/main/datasets | 10 | 25 GB |
|  | webKnossos | https://zarr.webknossos.org | 69 | 70TB |
| **In-progress** | Brain Image Library | https://www.brainimagelibrary.org/ |  |  |
|  | HuBMAP | https://portal.hubmapconsortium.org/ |  |  |
|  | OpenOrganelle | https://openorganelle.janelia.org/datasets |  |  |

### Amazon S3

Some of the most visible uses of OME-Zarr are part of the “Public Data” programs provided by large, commercial vendors like Amazon^47^ and Google^48^ to share community-critical datasets. Submissions to these programs are reviewed for overall value to the community, but if accepted, represent a particularly accessible resource.

#### Cell Painting Gallery

The Cell Painting Gallery^49^ contains Cell Painting (Bray et al. 2016) images and image-based profiles from many publicly available datasets, hosted by AWS Open Data Registry. Currently, the LINCS Dataset (Way et al. 2022) is available in the Cell Painting Gallery in OME-Zarr format. In LINCS, 110,012,425 A549 human lung carcinoma cells across 136 plates were treated with 1,571 compounds across 6 dose points. Morphology was captured by a standard Cell Painting workflow of five fluorescent channels covering eight organelles. Image data was converted to OME-Zarr using bioformats2raw with the Distributed-OMEZarrCreator wrapper (Weisbart and Cimini 2022). 1,790 morphological measurements were taken using CellProfiler (Kamentsky et al. 2011) which are also available in the Cell Painting Gallery. More Cell Painting datasets in the Cell Painting Gallery are planned for both conversion to OME-Zarr and browsability through IDR.

#### DANDI

DANDI, “Distributed Archives for Neurophysiology Data Integration” is supported by the US BRAIN Initiative as an open-access data repository for publishing and sharing neurophysiology data including electrophysiology, optophysiology, and behavioral time-series, as well as images from immunostaining experiments. As datasets get larger, downloading whole datasets for analysis or visualization becomes increasingly impractical. Thus, DANDI allows data to be operated on in the cloud using standardized methods and computational servers near the data. It enforces the organized structure of the Brain Imaging Data Structure (BIDS) and its microscopy extensions (Bourget et al. 2022), with the latest specification^50^ allowing such data to be stored in the OME-Zarr format. DANDI uses standardized metadata (JSON, JSON-LD), data organization (BIDS, or BIDS-like subset), and data storage formats (e.g., OME-Zarr) that allow the metadata to be queried before data access, and data to be accessed partially and at different resolutions. The archive stores open and embargoed data on Amazon S3 in the US and is supported by the Amazon Open Data program. The DANDI API has full programmatic support for uploading and accessing Zarr objects, and researchers can use the DANDI command line interface tool to upload BIDS organized Zarr data to DANDI. All datasets are also available as DataLad (Halchenko et al. 2021) datasets from GitHub^51^ with individual Zarr objects provided as separate repositories^52^ and linked to original datasets as git submodules. Since data itself resides on the versioned S3 bucket, such DataLad datasets provide access to TBs of OME-Zarr data with unambiguous versioning.

DANDI allows any researcher to view or compute on the data. Lightly pre-processed data and metadata are being made available via DANDI for the BRAIN Initiative and other projects. Two such datasets were produced by the BRAIN Initiative Cell Census Network (biccn.org) comprising 370 TB of data. About 60% of a single human brain hemisphere is available in a dataset with vascular, neuronal, and cellular stains (290 TB; an example 3D slab shown in Fig. 13) with an additional 65TB from the brains of two other participants. These data can be visualized by anyone on the planet due to horizontal scalability provided by AWS S3, the use of a streaming data storage format (OME-Zarr), and interactive visualization tools such as Neuroglancer, ITK-VTK-Viewer, and others described earlier. For the OME-Zarr data, each such object provides an option to directly open the object in the ITK-VTK-Viewer or for checking using the OME-Zarr Validator. To optimize data transfer, DANDI recommends OME-Zarr lightsheet imaging data on human brains to be stored with chunk sizes of (1,1,128,128,128) and with lossless Blosc+Zstd compression.

**Fig. 13.**
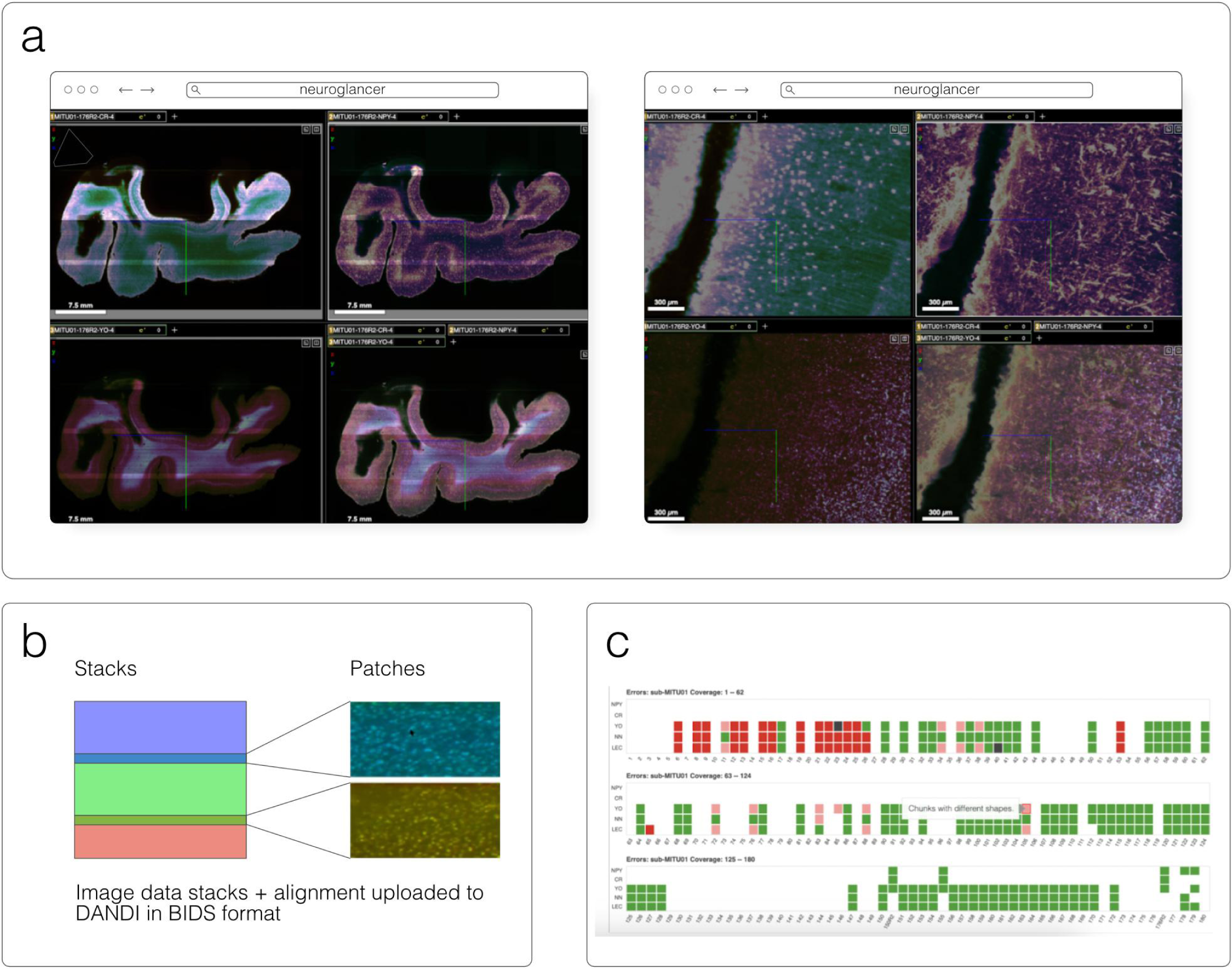
An OME-Zarr dataset on DANDI (ID: <u>DANDI:000108</u>) contains multiscale 5D datasets (time, channel, z, y, x), with metadata, scale, and transformations. A) DANDI:000108 includes multi-slab, multi-stain (NeuropeptideY-NPY, Calretinin-CR, YOYO1) data that are aligned and the coordinate transforms are stored in the Zarr files, allowing B) on-the-fly visualization and stitching of the slabs at multiple scales (two shown) using Neuroglancer. C) An HTML dashboard allows data submitter and any user on the Web to see the live status: samples+stains uploaded, their quality issues, and multiscale viewing of this 300+ TB dataset.

#### Others

A number of other projects like Allen Institute for Neural Dynamics (AIND)^53^ have also followed this strategy to achieve FAIR, Open and Reproducible Science. However, it is equally possible to host your own data on commercial services if funds are available. Glencoe Software, as the commercial arm of OME, regularly stores OME-Zarr data in buckets like s3://gs-public-zarr-archive^54^. A careful consideration is needed of the price per terabyte as well as egress costs incurred by users accessing such buckets.

#### Alternative S3 Providers

S3 has become a *de facto* standard API. As such, a number of providers offer commercial systems with many of the same features. Several publicly funded institutions dedicate themselves to providing large-scale infrastructure for sharing biological data and this is often achieved by providing “S3-like” storage.

#### BIA and IDR

An example of an institute providing a large-scale cloud repository is the European Bioinformatics Institute (EBI) who has done so by investing in object, or “cloud”, storage. The Image Data Resource (IDR) (Williams et al. 2017) and the BioImage Archive (BIA) (Hartley et al. 2021) are free public digital repositories for biological images hosted at EBI that are associated with peer-reviewed publications or are of value beyond a single experiment. The BIA strives to provide hosting for all such publications while the IDR focuses on a selection of reference datasets which are further curated. They accept submissions of many different image formats to enable easy and fast sharing. However, most of these formats have issues relating to metadata and to online visualization as discussed above.

The IDR’s need to store data on object storage led OME to begin discussions around a potential NGFF. As early adopters and implementers, the IDR found that such a file format alleviated many bioimaging bottlenecks at scale. Multi-terabyte 3D volumes (McDole et al. 2018) and 100 terabyte high-content cell painting screens (Bray et al. 2016), common to the IDR, can be more reasonably processed and accessed if converted in the repository to OME-Zarr. Additionally, use of OME-Zarr as a submission format reduced the burden on the curation team to deal with compatibility of custom formats.

In the BIA, OME-Zarr is chosen as its chunked architecture and available libraries and visualization tools enable on-the-fly visualization of the data, including 3D images, time series data and images with multiple channels. Moreover, it can accommodate extensive image level metadata, and has substantial community support.

To enable easier sharing and investigation of long-term solutions, images are converted from selected datasets, including most machine learning datasets, into OME-Zarr format. The converted representation is stored transiently on fast, S3 compatible storage. This enables serving the images for on-the-fly visualization in embedded viewers, visualization by desktop clients that can access OME-Zarr remotely such as napari or MoBIE, and provides fast access for on-the-fly analysis via Jupyter notebooks. The converted image representations are transient due to the need to balance resources between cheaper scalable storage for long term archival and fast storage for access to in-demand datasets. The OME-Zarr converted image representations are currently presented on separate websites, with plans to integrate these within the primary dataset pages. Images can be displayed using Vizarr, and an option to view them with the ITK viewer is also provided. As with all submissions, each image representation has a unique URI therefore the users can access and view them with their favorite online viewers, without the need to download. Already it is possible to submit datasets that have been converted into OME-Zarr and as this solution matures, submitters will be encouraged to pre-convert complicated datasets to lower the overhead on the curation teams.

#### Sanger

Data generated by Wellcome Sanger Institute and Newcastle University are made available on their public S3 buckets. That includes both array-based single cell sequencing data and microscope-based imaging data. The data conversion was performed via a bespoke data ingestion pipeline which translates every object into OME-Zarr, chosen to be the default image format for all the data generated from spatial technologies such as Visium, in situ sequencing, RNAscope, Xenium and Merfish.

#### SSBD

SSBD is a platform for Systems Science of Biological Dynamics to share and reuse bioimaging data hosted at RIKEN, Japan (Tohsato et al. 2016). It consists of two systems; SSBD:database for sharing highly reusable data with rich metadata and SSBD:repository for rapid sharing of all bioimaging data in journal papers. To support the development and validation of software tools for NGFF, we converted selected datasets in SSBD:database into OME-Zarr format using the bioformats2raw converter. The datasets include those obtained with light sheet fluorescence microscopy, STED super-resolution microscopy, FIB-SEM, and other state-of-the-art microscopies. They are currently stored and available on S3-compatible storage in the RIKEN HOKUSAI SS system.

#### On-premise

Even beyond conventional object storage, OME-Zarr provides a mechanism for consistently sharing bioimaging data. This can be as simple as putting the files on an existing web server or dedicated servers can be installed locally.

#### Zebrahub

Zebrahub^55^ is a comprehensive atlas of zebrafish development (Lange et al. 2023) paired with a fully interactive website to explore both single-embryo single-cell RNA sequencing datasets and light-sheet microscopy time-lapses. Light-sheet microscopy is a powerful imaging technique that, when combined with transgenic zebrafish expressing fluorescent proteins, can reconstruct the entire embryonic development at a cellular level. This results in terabyte-scale time-lapse datasets that contain 3D images acquired at regular intervals, often every 30 seconds. Acquiring multiple channels is necessary to answer specific biological questions, which further increases the data’s size and dimensionality. Storing, processing, visualizing, and disseminating such data becomes challenging. The OME-Zarr format addresses several of these issues by providing a well-established and maturing standard storage format supported by various image processing and visualization tools. The Zebrahub project and its accompanying website leverage OME-Zarr by providing image data downloads in OME-Zarr format and neuroglancer instances that allow for the interactive exploration of these image datasets (see Fig. 14). Sharing terabyte-scale high-resolution light-sheet imaging time-lapse datasets in a web browser via OME-Zarr is a significant contribution towards advancing observation-based discovery in developmental biology.

**Fig 14.**
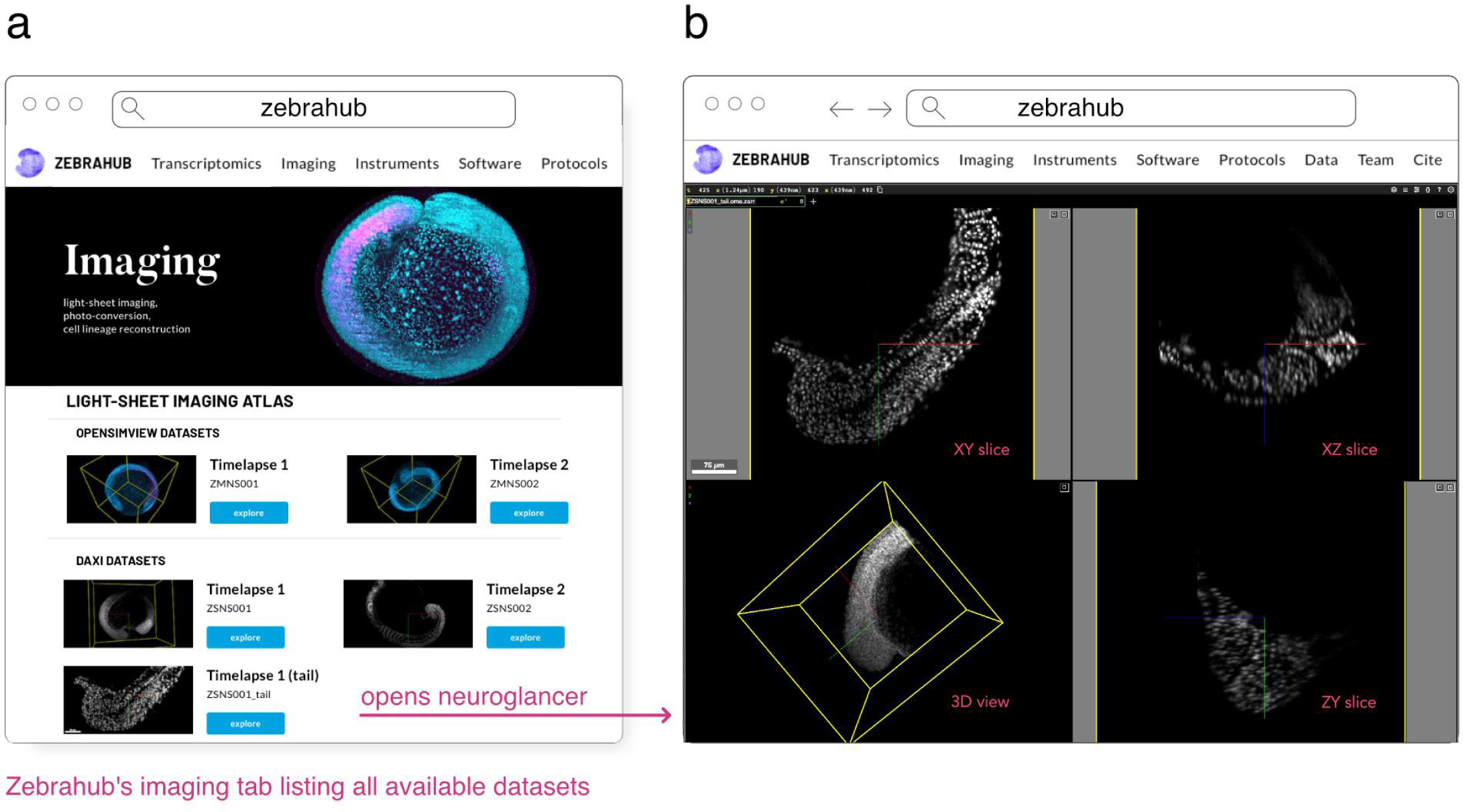
(a) ‘Imaging’ tab of the zebrahub.org website showing the list of imaging datasets made available. (b) By clicking on a particular dataset, the user is directed to a neuroglancer instance that allows interactive exploration of the dataset.

#### Hosting your own

Alternatively there are existing software solutions that expose an object-storage interface to files on disk like Minio^56^. The European Molecular Biology Laboratory in Heidelberg is currently hosting all of the reference data for the development of the MoBIE tool by running Minio on their own hardware. Examples of datasets formatted in OME-Zarr include https://github.com/mobie/covid-if-project, https://github.com/mobie/htm-test and https://github.com/mobie/spatial-transcriptomics-example-project. In the README of those repositories, a list of all S3 OME-Zarr URLs can be found which can be opened with viewers other than MoBIE. Sample data for the SpatialData library can also be found on EMBL’s S3 resource. A list is available under https://github.com/scverse/spatialdata-notebooks/tree/main/datasets.

Alternatively, dedicated, bioimaging server solutions are available for hosting and serving OME-Zarr data. As mentioned above, webKnossos makes all datasets accessible in the OME-Zarr format. To import data into webKnossos a conversion is required, whether automatically on browser upload or manual through filesystem import to OME-Zarr or WKW, the “webKnossos Wrapper” format. Once imported into webKnossos, datasets can be made publicly available via the dataset settings. This is an easy way to publish datasets directly from an institutional HPC storage. A collection of published OME-Zarr data is available at https://zarr.webknossos.org/.

Another possible mechanism for hosting OME-Zarr data is OMERO (Allan et al. 2012). The OMEZarrReader plugin to Bio-Formats is included in OMERO since version 5.6.5, enabling import of OME-Zarr data into any OMERO including the IDR. These are stored in OMERO’s internal repository with other file types and appear as images, or in the case of HCS data plates, in the standard hierarchy. Users can then use the OMERO API or clients (OMERO.figure, QuPath, or even R) to access the data. To enable OME-Zarr access to end users, services are available which expose an OME-Zarr compatible endpoint to clients.

#### In-progress

Finally, several repositories have plans of migrating all of their existing datasets to OME-Zarr as a common representation for dissemination.

#### Brain Image Library

Institutes with large scale parallel (HPC) file systems like the Pittsburgh Supercomputing Center, home to the Brain Image Library^57^ (BIL), are building tools to share OME-Zarr formatted data without object storage and without modifying the existing data repository^58^. Still in early development, these services will make 1000s of whole brain microscopy datasets, including murine, NHP and human, some expected to be over a petabyte in size, from the BICCN (BRAIN Initiative Cell Census Network) (BICCN Data Ecosystem Collaboration et al. 2022) and BICAN (BRAIN Initiative Cell Atlas Network) available in OME-Zarr where appropriate starting this year. To achieve this, microservices will automatically provide an OME-Zarr representation of the day on the fly to prevent data duplication. These microservices will enable all of the visualization software described above (Fig. 15) to make use of the data while permitting the high-performance computing facilities to optimize for their storage requirements. Similar APIs will be employed locally at BIL to allow HPC computational tools to interface with the data in a standard way, and with community derived tools developed around OME-Zarr.

**Fig. 15.**
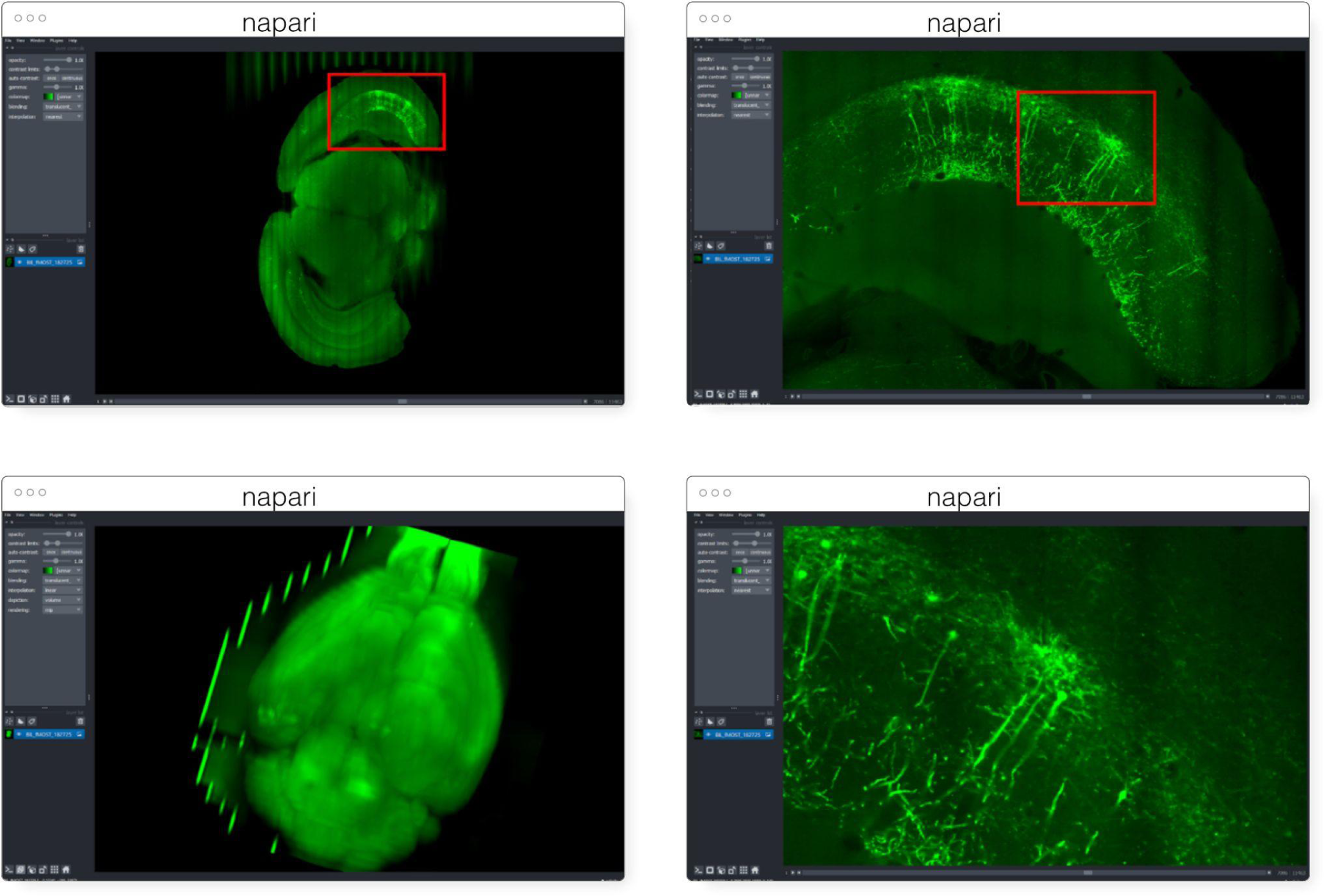
Multiscale OME-Zarr representation of a whole brain dataset^59^ (∼6TB compressed) archived at BIL and visualized over the internet in napari via the napari-ome-zarr plugin. Data is stored at BIL in an alternative format and is dynamically converted to OME-Zarr chunk-by-chunk and delivered to clients upon request.

#### HuBMAP

The HuBMAP (“Human Biomolecular Atlas Program”) consortium (HuBMAP Consortium 2019) adopted open-standard imaging formats and developed novel web-based technologies in order to simplify the server-side infrastructure traditionally required to support visualization of such a bioimaging resource. Consideration of infrastructure costs and a commitment to FAIR data access principles led the consortium to adopt OME-TIFF as an internal standard. In conjunction, members of the consortium developed Viv (Manz et al. 2022) – a client-side bioimaging visualization library – to remove a dependency on server-side rendering and enable flexible browsing of multi-terabyte datasets directly within the HuBMAP data portal website. Currently, HuBMAP is working to adopt OME-Zarr in order to support a wider diversity of modalities (e.g., large 3D volumes) as well as integrate additional data types (e.g., segmentations, ROIs, 3D rendering, high-content screening data, and non-image formats like AnnData) not supported by OME-TIFF. This data will be hosted on the Pittsburgh Supercomputing Center’s hardware.

#### OpenOrganelle

The OpenOrganelle (Heinrich et al. 2021) from the Janelia Research Campus hosts all of the datasets listed under https://openorganelle.janelia.org/datasets on AWS’s Open Data Registry. The data is visualizable in the browser using neuroglancer or via Fiji N5-plugins. The data is currently stored in a precursor to OME-Zarr known as the N5 format with “NGFF-compatible” metadata with conversion to OME-Zarr on the future roadmap.

## Discussion

Over the last twenty years, the number of bioimaging file formats has been a constant source of confusion and frustration. While that has often been a struggle that each user manages in isolation, increasingly data sizes from more sophisticated hardware and more advanced modalities are leaving users with significant infrastructure burdens for efficiently converting and sharing their imaging data. Data acquisition systems often design formats specifically for writing data quickly to timely capture the scale and the breadth of modern experiments. The tension between the requirements of quickly *writing* and quickly *reading* bioimaging datasets force both data providers and consumers to be aware of the costs of converting, relinking, downsampling, or otherwise modifying datasets for reuse. A one-time conversion of such “write-optimized” data can lower the overhead of repeated analysis and visualization of the data, but requires a widely adopted target format. With proper support, a small suite of storage file formats like HDF5, TIFF and Zarr can cover the essential use cases for optimally achieving the community’s scientific goals as has been achieved by other projects (e.g., PDB (Berman et al. 2012) and NetCDF (Unidata Ltd 1973)).

The strategy outlined in this paper is to encourage community cooperation towards a common representation. Increased focus from the community of developers accelerates the features delivered to the user community. Increases in the expressive power of the format through specifications, the number and ability of available tools, and data publicly available for end users all motivate further developments in each of the other areas. In turn, this progress drives the ability of the bioimaging community to better enact the FAIR principles. The growth of OME-Zarr tools, resources, and specifications, however, should not be taken as a reason to wait on adoption. The opposite is true. The hope is that more users and more developers will drive further growth, further unifying the bioimaging ecosystem. Users should identify whether or not the advancements detailed here will simplify and accelerate their scientific practice, and if so, are encouraged to start using OME-Zarr today. The community is growing and membership is open, free and encouraged. User feedback is critical to help make the most FAIR representation of bioimaging data possible.

## Code availability

All code described is available publicly under free and open-source licenses.

## Data availability

All data shown in the figures is available publicly under permissive licenses.

## Statements and Declarations

### Author Contributions

**Conceptualization:** Josh Moore; **Software:** Daniela Basurto-Lozada, Sébastien Besson, John Bogovic, Jordão Bragantini, Eva M. Brown, Jean-Marie Burel, Gustavo de Medeiros, David Gault, Satrajit S. Ghosh, Ilan Gold, Yaroslav O. Halchenko, Matthew Hartley, Dave Horsfall, Nathan Hotaling, Mark S. Keller, Gabor Kovacs, Aybüke Küpcü Yoldaş, Koji Kyoda, Merlin Lange, Tong Li, Prisca Liberali, Dominik Lindner, Melissa Linkert, Joel Lüthi, Jeremy Maitin-Shepard, Trevor Manz, Luca Marconato, Matthew McCormick, Khaled Mohamed, Josh Moore, William Moore, Wei Ouyang, Bugra Özdemir, Giovanni Palla, Constantin Pape, Lucas Pelkmans, Tobias Pietzsch, Stephan Preibisch, Norman Rzepka, Stephan Saalfeld, Sameeul Samee, Nicholas Schaub, Hythem Sidky, Ahmet Can Solak, David R. Stirling, Jonathan Striebel, Christian Tischer, Daniel Toloudis, Isaac Virshup, Alan M. Watson, Erin Weisbart, Kevin A. Yamauchi; **Resources:** Sébastien Besson, Matthew Hartley, Koji Kyoda, Shuichi Onami, Martin Prete; **Data curation:** Aybüke Küpcü Yoldaş, Frances Wong; **Visualization:** Erin E. Diel, Satrajit S. Ghosh, Gabor Kovacs, Albane le Tournoulx de la Villegeorges, Joel Lüthi, Matthew McCormick, Josh Moore, William Moore, Martin Prete, Christian Tischer, Daniel Toloudis, Alan M. Watson; **Writing - original draft preparation:** Daniela Basurto-Lozada, Sébastien Besson, John Bogovic, Eva M. Brown, Jean-Marie Burel, Gustavo de Medeiros, Erin E. Diel, Satrajit S. Ghosh, Ilan Gold, Yaroslav O. Halchenko, Matthew Hartley, Dave Horsfall, Mark S. Keller, Gabor Kovacs, Aybüke Küpcü Yoldaş, Tong Li, Prisca Liberali, Joel Lüthi, Jeremy Maitin-Shepard, Trevor Manz, Matthew McCormick, Josh Moore, Bugra Özdemir, Constantin Pape, Lucas Pelkmans, Tobias Pietzsch, Stephan Preibisch, Norman Rzepka, Christian Tischer, Daniel Toloudis, Petr Walczysko, Alan M. Watson, Frances Wong, Kevin A. Yamauchi; **Writing - review and editing:** Jean-Marie Burel, Xavier Casas Moreno, Beth A. Cimini, Yaroslav O. Halchenko, Nathan Hotaling, Mark S. Keller, Trevor Manz, Josh Moore, William Moore, Nils Norlin, Stephan Preibisch, Norman Rzepka, Stephan Saalfeld, Jonathan Striebel, Jason R. Swedlow, Erin Weisbart; **Supervision:** Omer Bayraktar, Beth A. Cimini, Nils Gehlenborg, Muzlifah Haniffa, Nathan Hotaling, Shuichi Onami, Loic A. Royer, Stephan Saalfeld, Oliver Stegle, Jason R. Swedlow, Fabian J. Theis

### Competing Interests

S.B., E.D., M.L., D.R.S. and J.R.S. are affiliated with Glencoe Software, a commercial company that builds, delivers, supports and integrates image data management systems across academic, biotech and pharmaceutical industries; J.M. and W.M. also hold equity in Glencoe Software. M.M. is affiliated with Kitware, Inc., a commercial company built around open-source platforms that provides advanced technical computing, state-of-the-art AI, and tailored software solutions to academic, government, and industrial customers. A.V., J.S. and N.R. are affiliated with Scalable Minds, a commercial company that builds, delivers, supports and integrates image analysis solutions. F.J.T. consults for Immunai Inc., Singularity Bio B.V., CytoReason Ltd, Cellarity, and Omniscope Ltd, and has ownership interest in Dermagnostix GmbH and Cellarity. N.A.H. and N.J.S. are contractors who work for Axle Research and Technology. S.B.S. and H.S. are affiliated with Axle Research and Technology and are contracted to the National Center for Advancing Translational Science, NIH. The remaining authors declare that they have no known competing financial interests or personal relationships that could have appeared to influence the work reported.

### Footnotes

1 https://zarr.dev/

2 https://nasa.github.io/Transform-to-Open-Science/year-of-open-science/

3 https://ngff.openmicroscopy.org/0.4

4 https://ngff.openmicroscopy.org/0.4/#hcs-layout

5 https://vitessce.io

6 https://www.allencell.org/pathtrace-rendering.html

7 https://mobie.github.io/

8 https://github.com/bigdataviewer

9 https://github.com/saalfeldlab/n5

10 https://napari.org/

11 https://github.com/ome/napari-ome-zarr

12 https://napari.org/stable/naps/4-async-slicing.html

13 https://github.com/scverse/napari-spatialdata

14 https://github.com/google/neuroglancer

15 https://github.com/ome/ome-ngff-validator

16 e.g. https://ome.github.io/ome-ngff-validator/?source=https://uk1s3.embassy.ebi.ac.uk/idr/zarr/v0.4/idr0062A/6001240.zarr

17 https://github.com/hms-dbmi/viv

18 https://github.com/hms-dbmi/vizarr

19 https://www.openmicroscopy.org/2020/12/01/zarr-hcs.html

20 https://webknossos.org/

21 https://cfe.allencell.org

22 https://allen-cell-animated.github.io/website-3d-cell-viewer/

23 https://pypi.org/project/nbvv/

24 https://github.com/gzuidhof/zarr.js/

25 https://github.com/mobie/bigdataviewer-n5

26 https://github.com/PreibischLab/BigStitcher-Spark

27 https://github.com/bigdataviewer/bigdataviewer-omezarr

28 https://ome-zarr.readthedocs.io/

29 https://fractal-analytics-platform.github.io

30 https://slurm.schedmd.com/documentation.html

31 https://www.youtube.com/watch?v=DfhRF1OW5CE

32 https://github.com/PolusAI/bfio

33 https://spatialdata.scverse.org/en/latest/

34 https://google.github.io/tensorstore/

35 https://github.com/PolusAI/nyxus

36 https://github.com/ome/ZarrReader

37 https://github.com/saalfeldlab/n5-ij

38 https://github.com/saalfeldlab/n5-viewer

39 https://idr.openmicroscopy.org/webclient/?show=project-502

40 https://www.dask.org/

41 https://colab.research.google.com/

42 https://mybinder.org/

43 https://github.com/glencoesoftware/bioformats2raw

44 https://www.glencoesoftware.com/products/ngff-converter/

45 https://imswitch.readthedocs.io/en/stable/

46 https://github.com/fsspec/kerchunk

47 https://aws.amazon.com/opendata/open-data-sponsorship-program/

48 https://cloud.google.com/storage/docs/public-datasets

49 https://registry.opendata.aws/cellpainting-gallery

50 https://bids-specification.readthedocs.io/en/stable/04-modality-specific-files/10-microscopy.html#file-formats

51 https://github.com/dandisets

52 https://github.com/dandizarrs

53 https://registry.opendata.aws/allen-nd-open-data/

54 https://glencoesoftware.com/ngff

55 http://zebrahub.org/

56 https://min.io

57 https://www.brainimagelibrary.org/

58 https://github.com/CBI-PITT/BrAinPI

59 https://download.brainimagelibrary.org/2b/da/2bdaf9e66a246844/mouseID_405429-182725/

